# A Digital Microfluidic Platform for the Microscale Production of Functional Immune Cell Therapies

**DOI:** 10.1101/2024.09.03.611092

**Authors:** Samuel R. Little, Niloufar Rahbari, Mehri Hajiaghayi, Fatemeh Gholizadeh, Fanny-Mei Cloarec-Ung, Joel Phillips, Hugo Sinha, Alison Hirukawa, David J.H.F. Knapp, Peter J. Darlington, Steve C.C. Shih

## Abstract

Genetically engineering human immune cells has been shown to be an effective approach for developing novel cellular therapies to treat a wide range of diseases. To expand the scope of these cellular therapies while solving persistent challenges, extensive research and development is still required. Electroporation has recently emerged as one of the most popular techniques for inserting biological payloads into human immune cells to perform genetic engineering. However, several recent studies have reported that electroporation can negatively impact cell functionality. Additionally, the requirement to use large amounts of cells and expensive payloads to achieve efficient delivery can drive up the costs of development efforts. Here we use a digital microfluidic enabled electroporation system (referred to as triDrop) and compare them against two state-of-the-art commercially available systems for the engineering of human T cells. We describe the ability to use triDrop for highly viable, highly efficient transfection while using substantially fewer cells and payload. Subsequently, we perform transcriptomic analysis on cells engineered with each of the three systems and show that electroporation with triDrop lead to less dysregulation of several functionally relevant pathways. Finally, in a direct comparison of immunotherapeutic functionality, we show that T cells engineered with triDrop have an improved ability to mount an immune response when presented with tumor cells. These results show that the triDrop platform is uniquely suited to produce functionally engineered immune cells while also reducing the costs of cell engineering compared to other commercially available systems.

---

Reprogramming the functionality of human T cells by inserting novel biological payloads has shown to be a promising avenue of therapeutic development.^1^ Removing immune cells from a patient, functionally modifying the cells to fight disease, and reinjecting them into the patient has been shown as a viable treatment for both hematological cancers,^2–4^ and solid cancers.^5^ However, manufacturing of these therapies is challenging,^1^ and a drawback with current available therapies is the lack of specificity can cause deleterious side effects.^6^ Efforts have been made to engineer immune cells for improved targeting to avoid so called “on-target, off-tumour” toxicities.^7^ However, developing cellular therapies for cancer that is affordable, as well as safe and efficient will require additional complex genetic engineering and substantial R&D.^8,9^

A key challenge in the cell engineering pipeline is the delivery of biological payloads across the cell membrane and into the cells, which must be done efficiently while preserving the viability, and functionality of the cells. There are a handful of recently reported techniques that can introduce payloads into cells either biologically (retroviruses,^10^ lentiviruses,^11,12^ herpes simplex virus,^13^ adenovirus,^14^ and adeno-associated virus).^15^ chemically (cationic lipids,^16^ lipid nanoparticles),^17^ or physically (electroporation,^18^ mechanoporation,^19^ sonoporation,^20^ or microinjection).^21^ Concerns over immunogenicity, semi-random transgene integration, and cytotoxicity have resulted in viral transduction becoming less popular,^22,23^ while low transfection efficiencies have led to decreased enthusiasm for chemical techniques.^24^ Physical transfection techniques are becoming the preferred approach as they generate temporary nanopores in the cell membrane allowing the cargo suspended in the surrounding media to permeate the cell where it remains trapped after the pores heal.^25^ Given that pre-clinical R&D is a substantial driver of cost and time when bringing a new cell therapy to market,^26^ a platform capable of automating laborious procedures and processing numerous samples in parallel, while requiring only small inputs of cell and reagents per reaction could reduce the cost and length of many cellular therapy development programs. Currently, there are several popular commercially available platforms used for cell therapy, however these platforms either require performing each reaction serially (one-at-a-time) for testing multiple conditions,^27^ or require large cellular and reagent inputs,^28^ which is expensive and at times requires pooling numerous donors together convoluting the data.

Microfluidic-based platforms are emerging as potential technologies for the physical transfection of human immune cells using techniques such as mechanical squeezing or compression,^19,29–32^ fluidic shearing,^33,34^ and electroporation.^35,36^ A primary goal for this field has been to develop a platform capable of clinical-scale production, specifically, a device that is able to efficiently insert a single type of payload into cells while operating continuously with a throughput > 10^6^ cells / minute.^37^ Towards that goal, Weaver et al.^38^ recently detailed the final results of phase 1 clinical trial where a microfluidic mechanoporation method was used to produce four doses per patient each containing upwards of 5 x 10^6^ cells / kg for patients with HPV16+ solid tumours. Another example is the development of a viscoelastic-mechanoporation method from Sevenler et al.^34^ capable of processing 2.5 x 10^8^ cells / minute. These are important techniques for clinical manufacturing, but we are not aware of a robust, parallel microfluidics platform capable of inserting multiple unique payloads and conditions while minimizing the cost and consumption of each single reaction.

We previously published a novel three-droplet system that allowed for efficient electroporation (EP) of human T cells on a digital microfluidic (DMF) platform.^39^ Our technique minimizes the generation of electrical current and protects the cells from harmful electrochemical reactions that can occur during EP. Further, our proposed platform (referred to as triDrop) required < 50,000 cells per reaction and by using the multiplexing capabilities of DMF, numerous reactions can be performed in parallel. However, a major drawback of electroporation is that cells engineered via EP have been shown in the past to suffer from impaired functionality and genetic dysregulation post transfection.^37,40,41^ Here, we use our triDrop electroporation system and compare against two commercially available state-of-the-art electroporation systems: Invitrogen Neon Electroporation System^TM^ (hereby referred to as the Neon), and the Lonza 4D-Nucleofector^TM^ (hereby referred to as the Nucleofector). Herein we describe how triDrop efficiently transfects reduced amounts of cells while using less payload compared to the Neon and Nucleofector systems. Additionally, triDrop can deliver both simple (mRNA) and complex (CRISPR gene-editing reagents) payloads with improved viability and proliferation capabilities after EP. We conducted a transcriptomic analysis to benchmark triDrop, revealing that cells engineered with triDrop exhibit significantly less genetic dysregulation compared to using Neon and Nucleofector. Finally, we show a proof-of-concept immunotherapeutic assay, demonstrating that the triDrop platform is uniquely suited as a miniaturized platform for the research and development of novel immunotherapies.

## Results and Discussion

### Comparing electroporation platforms

We analyzed three electroporation (EP) systems that are schematically overviewed in **Figure 1a-c**. **Figure 1a** is the Invitrogen Neon Electroporation System^TM^ (Neon)^42^, the system operates by placing 10 µL (or 100 µL) of cells and payload suspended in EP buffer inside a capillary tube with an anode placed at the top of the capillary tube and the bottom submerged in an electrolytic buffer that is in contact with a cathode. Applying a voltage across the anode and the cathode generates an electric field throughout the capillary tube. **Figure 1b** is the Lonza 4D-Nucleofector and it operates by placing 20 µL of EP buffer containing cells and payload into a cuvette with parallel anode and cathode. The sample is placed directly between the electrodes and a voltage is applied. **Figure 1c** is a recently published,^39^ droplet-based, electroporation system that relies on digital microfluidics hereby referred to as the triDrop system. The triDrop operates by merging three 1 µL droplets into a sequential chain, the outer droplets are comprised of a high conductivity media (∼ 16 mS / cm) and the inner droplet is comprised of low conductivity buffer (∼ 8 mS / cm) and contains cells and payload. We previously demonstrated that by placing the anode and cathode in contact with the outer droplets and applying a voltage, a homogenous electric field is generated across the inner droplet inserting the payload into the cells while protecting the cells from excessive current generation and harmful electrolytic byproducts often found at the anode and the cathode during electroporation. The Neon and Nucleofector have been shown extensively for transfection of primary human T cells, however we propose that the triDrop can offer two key advantages over these existing systems. First, is the ability to achieve high performance transfection while using fewer cells and less payload providing up to a 20-fold reduction in the overall cost of T cell engineering (**Table S1**). In addition, using lower quantities of cells per reaction enables larger libraries to be tested on a single donor and opens the possibility to perform electroporation with rare cell types. Second, by limiting the exposure of cells to excessive electrical current and harmful electrolytic byproducts (such as pH changes as a result of the reduction and oxidation of water molecules, chlorine and hydrogen gas bubbles, and metal ions),^43^ the health and functionality of the cells can be preserved post-electroporation. It has been shown that joule heating as a result of electrical current, and exposure to electrolysis are significant contributing factors to cell death as a result of EP,^44^ and that cell death increases as the cells get closer to the anode and cathode^45^. The triDrop was specifically designed to isolate cells away from the anode and cathode, reducing exposure to the electrolytic byproducts, and to create a highly resistive environment limiting the generation of electrical current.^39^ Further, using fewer cells than the recommendation given by the Neon and Nucleofector would reduce the overall electrical resistance of their system, leading to increased electrical current during pulsing and potentially more joule heating and severe electrolysis.^46^

**Figure 1.**
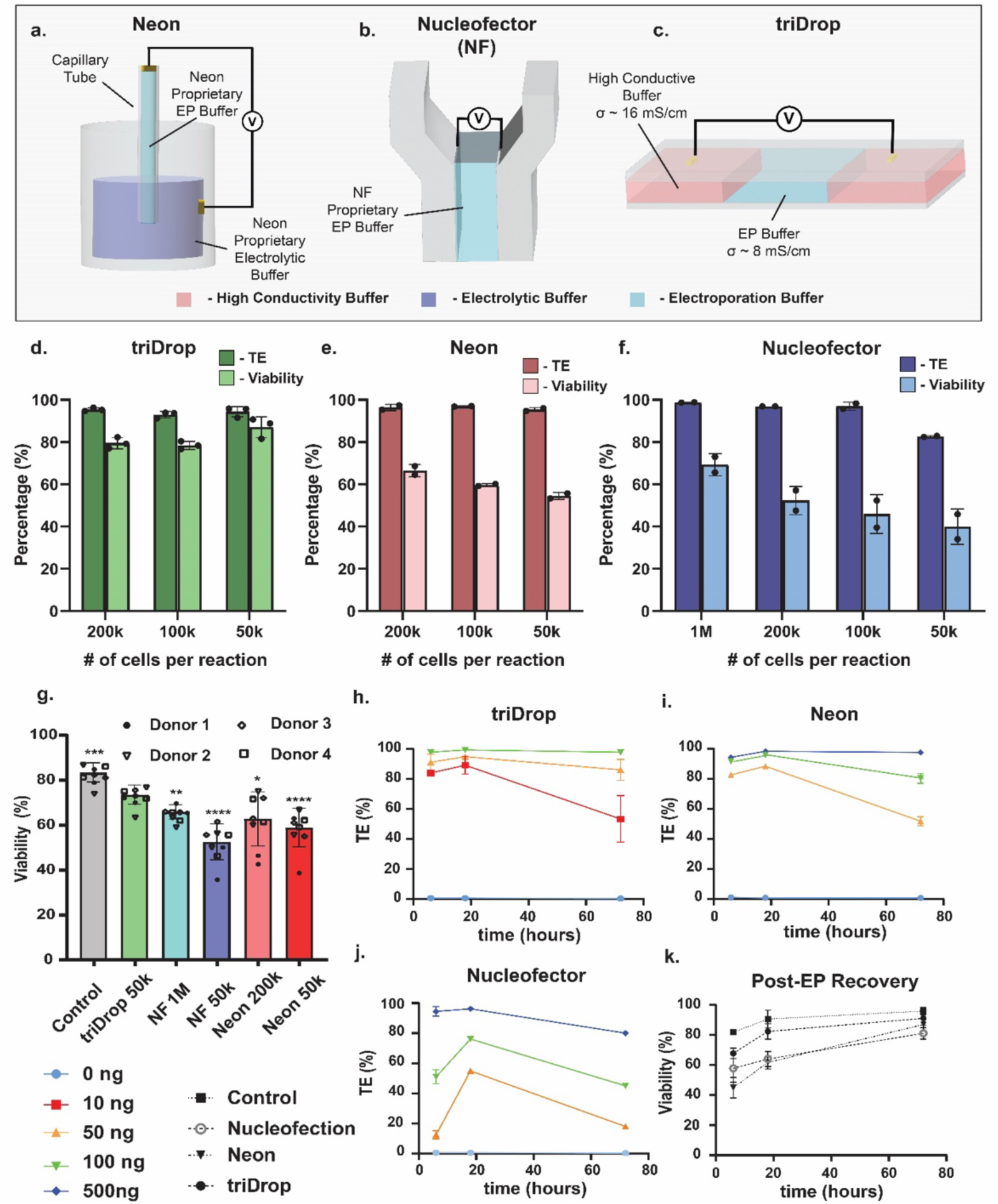
Three electroporation systems. Schematic overview of three electroporation systems used in this work where a) Neon, b) Nucleofector, and c) triDrop. Bar graphs depicting transfection efficiency (TE; dark colours) and viability (light colours) when performing electroporation using varying numbers of cells per reaction for d) triDrop, e) Neon, and f) Nucleofection. g) Viability measurements taken six hours post-EP for cells electroporated using either the manufacturer recommended number of cells or 0.5 x 10^5^ cells per reaction. Four donors (shown as polygons) were used for this study with two technical replicates. Statistical significance markers represent comparison between triDrop and all other conditions (n = 8). Line graphs depicting transfection efficiency over 72 hours post-EP when using 10 ng (red line), 50 ng (orange line), 100 ng (green line) and 500 ng (blue line) of mRNA for h) triDrop, i) Neon, j) Nucleofection. k) Line graph depicting viability over 72 hours post-EP for cells electroporated with all three EP systems and a control. All error bars represent mean ± 1 SD. Statistical n.s indicates no significant difference, *, **,***, and **** represent p-values below 0.05, 0.01, 0.001 and 0.0001 respectively. Statistical analysis was performed using a Student’s t-test.

We hypothesize that the triDrop platform has the ability to use fewer cells per reaction to achieve high transfection efficiency and cell viability post-electroporation. To test this hypothesis, three primary T-cell concentrations (0.5, 1, and 2 x 10^5^ cells per reaction) were electroporated with an eGFP-mRNA payload using one out of the three systems. As shown in **Figure 1d**, we found that while all three platforms were able to deliver mRNA molecules with efficiencies > 95 %, the triDrop had the highest reported viability as measured 18 hours post electroporation. The triDrop maintained viabilities above 80 % for 1, 2 x 10^5^ cell conditions and 85 % viability for electroporation reactions using 0.5 x 10^5^ cells. In comparison, **Figure 1e and 1f** show the Neon and Nucleofector achieved peak viabilities of ∼ 65 % (in line with previously published results^47^) when using the manufacturer recommended condition (1 x 10^6^ cells per reaction for Nucleofection, and 2 x 10^5^ cells per reaction for Neon). Interestingly, reducing the number of cells per reaction results in decreasing viabilities for both the Neon and the Nucleofector, achieving ∼ 60 % and ∼ 45 % viability respectively for EP reactions when using only 0.5 x 10^5^ cells per reaction (**Figure 1g**). To validate the effects of using reduced cell amounts on viability, we conducted EP using either the recommended number of cells or 0.5 x 10^5^ cells per reaction and measured viability 6 hours post-EP. While all systems led to a reduction in viability compared to the control, the triDrop system showed a significant improvement in viability in comparison to the Neon and Nucleofector regardless of how many cells were used. In addition, reducing the number of cells used by the Nucleofector leads to a significant reduction in viability (65 % vs. 51 %, P = 0.001), which could be related to our hypothesis of high current generation and electrolysis.

Next, we investigated the effects of using differing amounts of payload per reaction on transfection efficiency and viability. Cells were electroporated with a range of mRNA amounts using one of the three systems at their optimal cell concentrations (0.5 x 10^5^ cells for triDrop, 2 x 10^5^ cells for Neon, and 1 x 10^6^ cells for Nucleofector) and were analyzed with flow cytometry after 6, 18, and 72 hours. **Figure 1h-j** shows line graphs depicting transfection efficiency for each condition over 72 hours following EP. While the Neon and triDrop perform similarly, the Nucleofector requires approximately 10 times as much payload to achieve results comparable to the triDrop (500 ng vs 50 ng). In fact, the triDrop achieved > 80 % transfection efficiency while using the lowest tested amount of mRNA (10 ng). Furthermore, **Figure 1k** shows the measured viability of each system over 72 hours when using 100 ng of mRNA per reaction for the triDrop, 500 ng for Neon, and 1000 ng for Nucleofection (optimal amounts based on the results from **Figure 1h-j**). The data confirms that the triDrop has the highest viability of the three systems and by 18 hours post-EP cells treated with the triDrop have a viability of > 80 % compared to a viability of ∼ 60 % (consistent with results shown in **Figure 1d-f**). It is only after 72 hours post-electroporation that all three systems have a recovered viability > 80 %. When the cells are permeabilized by EP, the concentration gradient between the surrounding environment and the inside of the cell leads to payload being driven through the pores across the cell membrane.^48^ TriDrop uses smaller volumes for the system (electroporated volume of ∼ 1 µL), and as such, substantially less payload is required to achieve concentrations optimal for insertion into the cell.

### Knockout and knock-in gene-editing efficiency

The well-characterized CRISPR system has been used for engineering human immune cells to create new cellular immunotherapies.^28^ Additionally, screening large libraries of CRISPR edits has been shown as a successful avenue for immunotherapeutic discovery.^27^ The first step for such an application is to use delivery methods (such as electroporation) to introduce gene-editing components into cells. First, we investigated the performance of all three systems to conduct CRISPR knockouts targeting the TRAC locus. A range of cell concentrations were electroporated with an sgRNA cocktail complexed with a Cas9 protein to form a ribonucleic protein (RNP) and knockout efficiency was evaluated by staining with an anti-human alpha-beta TCR antibody and analyzed with flow cytometry four days post transfection. **Figure 2a-c** show the effects of knockout with all three systems using 0.5, 1, 2 x 10^5^ cells per reaction as well as a 1 million cell condition for the Nucleofector. In these experiments, RNP was normalized to the reaction volume to ensure consistent payload concentration as recommended by Hultquist et al.^49^ and Oh et al.^50^ and was added at a ratio of 50 pmol per 20 µL of reaction.. The triDrop achieved an optimal knockout efficiency of 75 % with a viability of 80 % when using 0.5 x 10^5^ cells per reaction, and the Neon and Nucleofector were both able to achieve knockout efficiencies ∼ 95 % when using higher numbers of cells. However, decreasing cell amounts to 0.5 x 10^5^ cell per reaction led to the efficiency dropping to 78 % and 70 % respectively.

**Figure 2.**
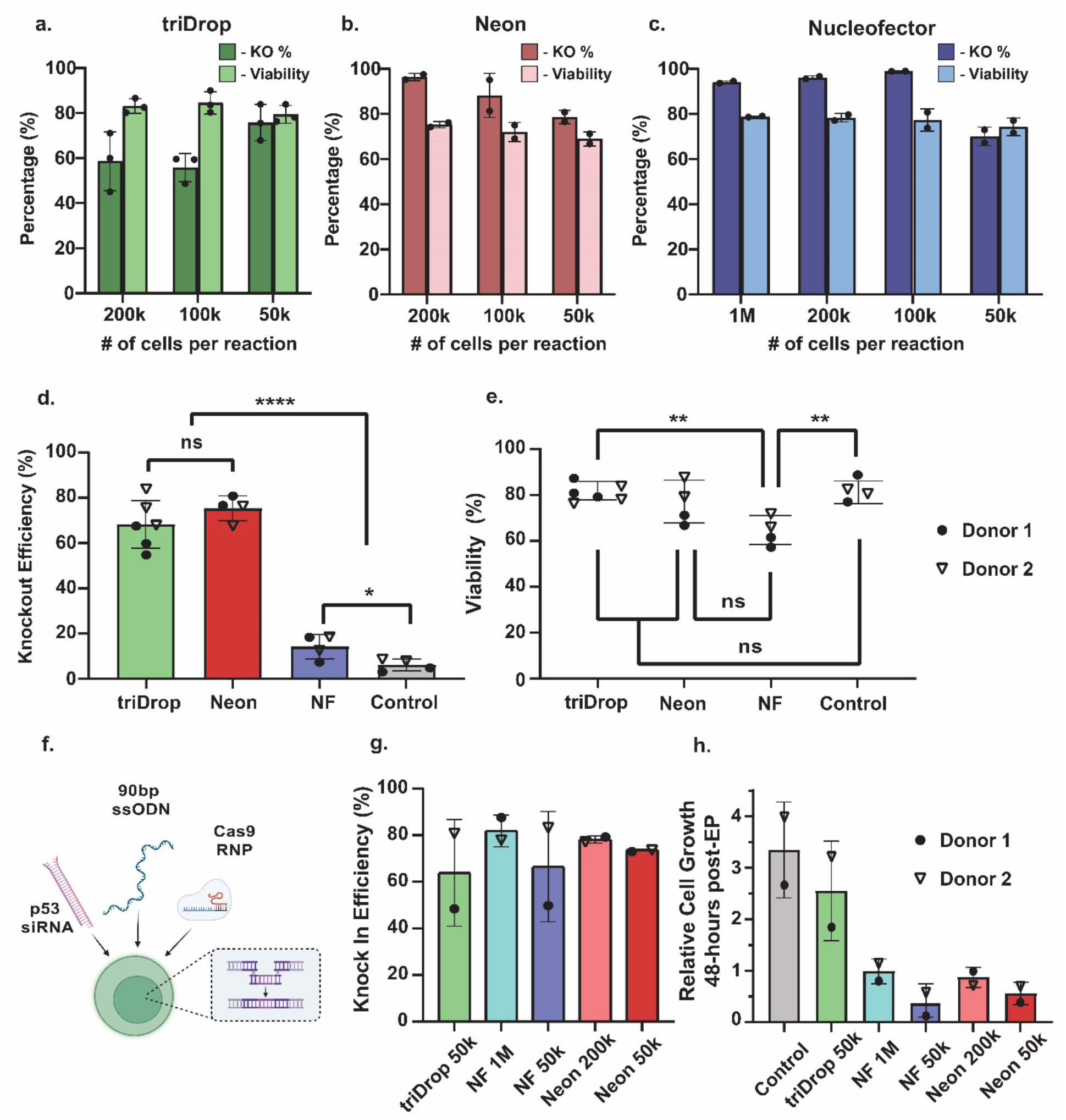
Gene editing. Bar graphs depicting knockout efficiency (dark colours) and viability (light colours) measured four days post EP when electroporating varying amounts of cells per reaction when using a) triDrop, b) Neon, and c) Nucleofector. Bar graphs showing d) knockout efficiency and e) viability for 0.5 x 10^5^ cells electroporated with 2.5 pmol of RNP using the three EP systems (n = 4 - 6). f) Schematic showing the payloads required for CRISPR knock-in (created with BioRender.com). Bar graphs showing g) knock-in efficiency and h) relative cell growth post EP for two donors electroporated using manufacturer recommended number of cells or 0.5 x 10^5^ cells per reaction. All error bars represent mean +/- 1 SD. Statistical n.s indicated no significant difference, *, **,***, and **** represent p-values below 0.05, 0.01, 0.001 and 0.0001 respectively. Statistical analysis was performed using a Student’s t-test.

Given the substantial volume differences between the three systems, adding payload in proportion to volume leads to a significant difference in the total amounts of Cas9 enzymes and sgRNA needed per reaction (i.e. 50 pmol of RNP for 1 nucleofection reaction vs 2.5 pmol for 1 triDrop reaction). To account for this difference, we tested conditions by normalizing payload to the number of cells being electroporated via addition of 50 pmol of RNP for every 1 million cells being used as recommended by Roth et al.^28^ We found that when using the Nucleofector, decreasing the payload in proportion to the number of cells being electroporated led to a decrease in knockout efficiency suggesting that volumetric normalization is more important than cellular normalization for achieving high delivery efficiency (**Figure S1**). **Figure 2d and e**, show a side-by-side comparisons of two donors when electroporating only 0.5 x 10^5^ cells with 2.5 pmol of Cas9 RNP. Both the Neon and the triDrop were able to achieve 75 % and 68 % knockout efficiency respectively with no significant difference between the two (p = 0.26), however, the Nucleofector was only able to achieve 14 % knock out efficiency with these conditions, which is significantly lower than the other two systems (p < 0.0001). Additionally, we observed that the decreased number of cells had a significant impact on the health of cells electroporated with the Nucleofector even after four days of recovery. Compared to the control the triDrop and Neon, both had no significant differences in cell viability (p = 0.67, and p = 0.3 respectively) whereas the Nucleofector had viabilities significantly less than both the control (p = 0.004) and cells treated with the triDrop (p = 0.0008). These show further evidence that the triDrop system can achieve efficient editing while requiring significantly less expensive payload.

After validating triDrop as a reliable method for knockout, we introduced donor templates to perform CRISPR knock-ins using the three methods. We followed a recent protocol published by Cloarec-Ung et al.^51^, which involved simultaneously co-delivering a Cas9 RNP targeting the SRSF2 gene, a 90 bp ssODN HDR-template, and an siRNA molecule targeting p53 (**Figure 2f**; all sequences in **Table S2**). Cells from two donors were electroporated using the Neon with 0.5 and 2 x 10^5^ cells per reaction, the Nucleofector using either 0.5 or 10 x 10^5^ cells per reaction, or the triDrop with 0.5 x 10^5^ cells per reaction. The amount of payload was normalized to the reaction volume such that the Nucleofector and Neon used 20- and 10-times more payload than the triDrop respectively. Cells were collected 48 hours post EP and the total number of viable cells were counted prior to being lysed and prepared for knock-in confirmation via Sanger sequencing. **Figure 2g** shows that all systems achieve average knock-in efficiencies of > 60 % and up to 80 % insertion efficiency, which is in line with recently published results.^52^ Interestingly, as shown in **Figure 2h**, we observed that the cells electroporated with the triDrop had proliferation capacities most similar to the control (2.5 fold increase in total viable cells vs 3.3 fold increase respectively), and more surprisingly, Nucleofector and Neon showed minimal proliferation post-EP. When using only 0.5 x 10^5^ cells per reaction, these systems showed a reduction in proliferative capacity, a 0.3 and 0.5 fold-change respectively in total viable cells demonstrating that using fewer cells with these commercial systems leads to impaired proliferation capabilities. These data suggest that all three systems can transfect both easy-to-deliver (mRNA) and hard-to-deliver payloads (Cas9 RNPs + HDR templates). However, the triDrop offers substantial improvements in both cell viability and proliferation capacity immediately after electroporation as well as requiring significantly less payload (at least 10x) and fewer cells.

### Transcriptomic analysis

The triDrop was developed to perform the efficient electroporation of primary human T cells for the purposes of developing and testing cellular therapies. To this end, it is important that cells engineered with the triDrop maintain their functionality following electroporation. To test for these effects, we carried out transcriptional profiling of human T cells post-electroporation. In the first type of test, we used qPCR to look at critical cytokines (IL-2, TNF-α, and IFN-γ) in the immune system that play significant roles in cellular therapies.^53^ Dysregulation of these genes lead to non-specific response or an impaired response in the presence of a target antigen.^54,55^ Cells were electroporated in all three systems using either manufacturer recommended conditions or 0.5 x 10^5^ cells per reaction and RNA was recovered from the cells 6 hours post EP. As shown in **Figure 3a-c**, cells treated with the triDrop shows no significant dysregulation in any of the three examined genes. Cells treated with the Neon, regardless of the number of cells used, shows no significant dysregulation of TNF-α, and IFN-γ, but there is a significant upregulation of IL-2 (relative fold change of 4.99, P = 0.005, and 5.11 P = 0.003) compared to the control. Similarly, when using the Nucleofector, cells treated with the manufacturer recommended conditions showed no dysregulation of the three genes, however, when using 0.5 x 10^5^ cells per EP reaction, IFN-γ was significantly downregulated (relative fold change of 0.38, P = 0.0006), indicating that cells electroporated under this condition may experience a reduced capacity to secrete this important cytokine. Intrigued by the results above, we used RNA sequencing to examine the entire transcriptional landscape of cells treated with all three systems. Electroporated cells from three different donors using the manufacturer recommended conditions for the Neon and Nucleofector and using 0.5 x 10^5^ cells per reaction for the triDrop were used to study the differential expression of genes between treated samples and a control. **Figures 4a-c** show volcano plots depicting the differential gene expression for all three electroporation systems compared to the control (with no electroporation). The x-axis depicts the log fold change of expression with the left and right displaying down- or up-regulated expression respectively, and the y-axis depicting the confidence in the gene expression changes represented by -log10(p-values). Using the cut-off metrics of > 1 or < −1 log fold change and a p–value of 0.05, we sorted genes into the categories of significantly dysregulated (red dots) or non-significant genes (grey dots). In addition, genes meeting the less stringent condition of p–values < 0.1 are shown as yellow dots. Based on these cutoffs, the Neon shows a dysregulation of 105 genes, the Nucleofector showing 89 genes, and only 32 genes for the triDrop. A principal component analysis showed the triDrop to be most similar to control cells (**Figure S2**). To understand these results, we classified them by grouping the genes that are commonly dysregulated between the systems **(Figure 4d)**. Six genes are commonly dysregulated between all systems and are shown in **Table S3.** Interestingly, 34 genes are commonly dysregulated between the Neon and the Nucleofector but not dysregulated with the triDrop. A collection of four such genes are highlighted in **Figure 4e-h**. The first two genes, PPP1R15A (2.7- and 2.8-fold increase for Neon and Nucleofector respectively), and SESN2 (3.6- and 2.9-fold increase for Neon and Nucleofector respectively), are implicated in integrated stress response pathways with SESN2 being implicated in pathways responding to oxidative DNA damage^56^ and PPP1R15A encoding for the growth arrest and DNA damage inducible protein.^57^ The upregulation of these two genes may partially explain the reduced viability and proliferation seen by cells electroporated with the Neon and the Nucleofector. The two other genes are TSC22D3 (2.9- and 2.4-fold change for Neon and Nucleofector respectively) and CD48 (−1.7- and −1.4-fold change for Neon and Nucleofector respectively) which are both known to affect the functionality of immune cells especially in the context of immunotherapy. First, TSC22D3 encodes for an anti-inflammatory molecule that is known to have immunosuppressive effects.^58^ The upregulation of this gene has been shown to compromise anti-tumour immunity in mice^59^ and corresponds with non-responsiveness in anti-CD19 CAR T trials in humans.^60^ Second, CD48 is a crucial member of lymphocyte activation and down regulation of this gene may impair the ability of T cells to respond to antigen presenting cells.^61^ Therefore, these two genes could impair the ability of cells engineered with the Neon and Nucleofector to adequately perform in immunotherapeutic assays.

**Figure 3.**
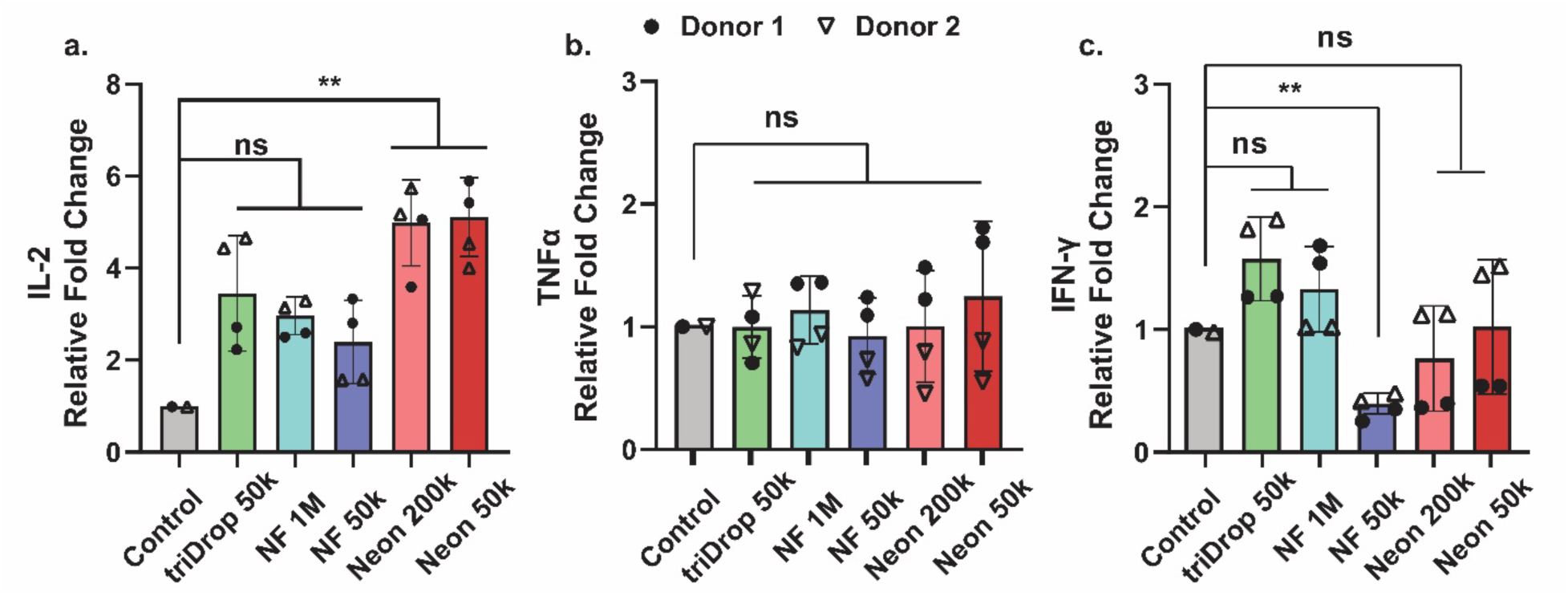
qPCR examination of key genes for cellular therapies. Bar graphs showing relative gene expression (2^−ΔΔc_t_^) 6 hours post EP for two donors electroporated using manufacturer recommended number of cells or 0.5 x 10^5^ cells per reaction for a) IL-2, b) TNF-α, and c) IFN-γ. All error bars represent mean +/- 1 SD. Significance n.s indicates no significant difference, and ** represents p-values below 0.01. Statistical analysis was performed using a Student’s t-test. (n = 4).

**Figure 4.**
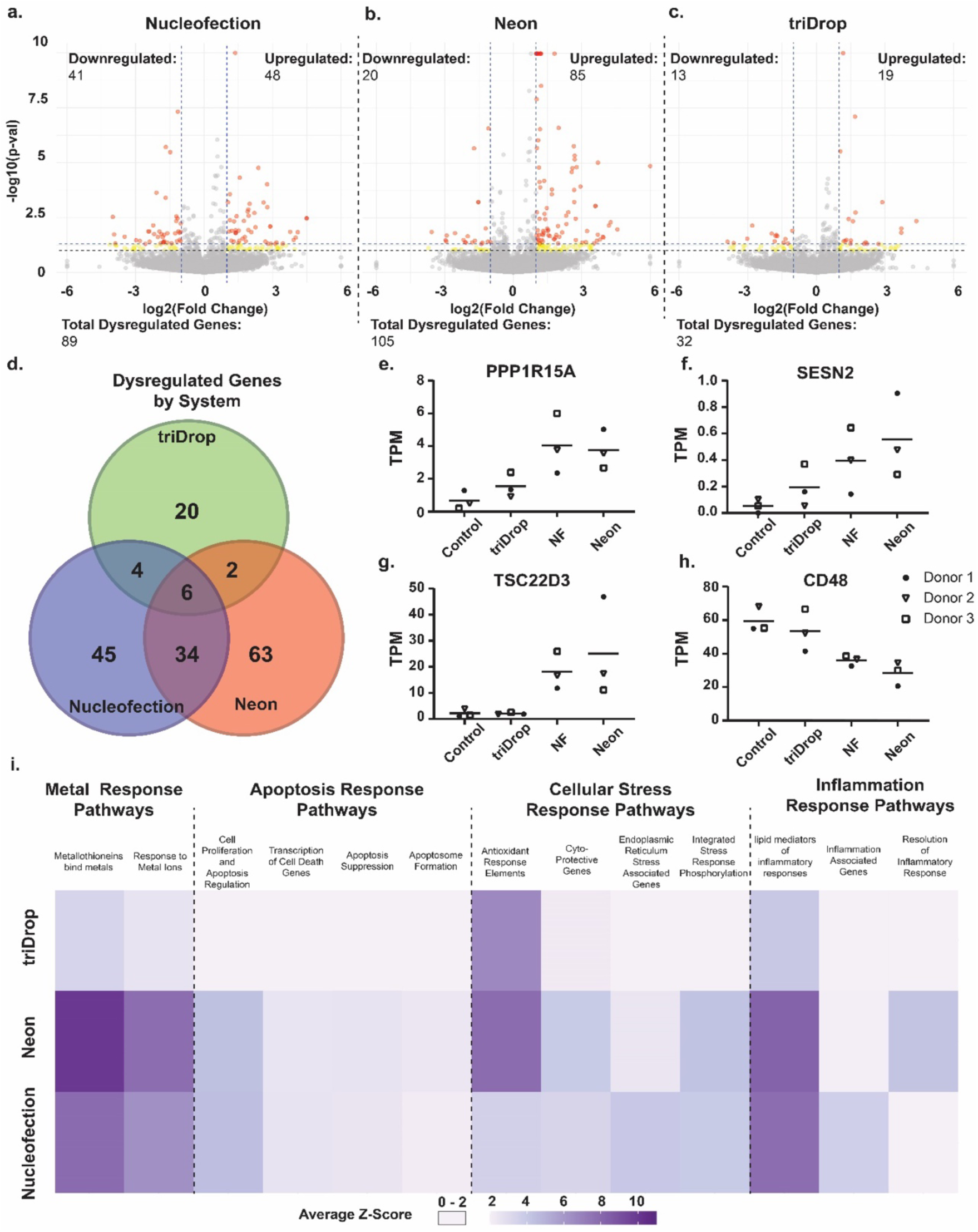
Transcriptomic analysis. Volcano plots depicting p-values and fold change for individual genes for cells from three different donors 6 hours post EP when electroporated with a) Nucleofection, b) Neon, and c) triDrop. Genes with a log fold change > 1 or < −1 with a p value < 0.05 (red dots) and < 0.1 (yellow dots) are highlighted. d) Venn diagram showing the number of genes that are uniquely dysregulated or mutually dysregulated between the three EP systems. Transcripts per million (TPM) values for three donors comparing the control, triDrop, Nucleofector, and Neon for e) PPP1R15A, f) SESN2, g) TSC22D3, and h) CD48. i) Heatmap showing the average Z-scores for 13 selected pathways for cells electroporated with all three EP systems.

For a more complete understanding of how genes were dysregulated in response to EP, all genes were assigned a z-score to quantify the different levels of expression relative to the control. Genes were then grouped into high level genetic pathways using the Reactome data base^62^ and the average z-score for the whole pathway was calculated. Out of the ∼ 2600 pathways analyzed, the triDrop showed a dysregulation of 79 pathways (defined as a pathway with an average z-score > 2.0 or < −2.0), the Neon with 130, and the Nucleofector showing 134. **Figure 4i** summarizes a collection of noteworthy pathways that are further documented in **Table S4.** As shown, all systems upregulate pathways responding to metal ion contaminates with the lowest z-score of upregulation shown by the triDrop. It is known that metal particulates can be secreted from the anode and cathode during electrolysis.^43^ The difference in z-scores observed for these pathway may be due to the Neon and Nucleofector placing cells in direct contact with at least one electrode whereas the triDrop isolates cells from both limiting contact with metal contaminates. Additionally, the Neon and the Nucleofector upregulate pathways corresponding to apoptosis and cellular stress which may further explain earlier data showing reduced viabilities and proliferation capabilities for cells treated with these systems. Finally, we observed upregulation of three inflammation pathways which are minimally upregulated or not upregulated with the triDrop. Upregulation of these pathways is concerning for cells being engineered for immunotherapy applications because chronic inflammation is known to lead to premature T cell exhaustion^63^ which can impair the function of CAR T cells.^64^

In summary, it is evident that cells engineered with the triDrop system have fewer dysregulated genes and dysregulated pathways relative to the Neon and Nucleofection systems. Using the Neon or Nucleofector, up-regulates genes and pathways associated with cellular stress, impairment of anti-tumour activity, as well as inflammation, which are either not upregulated or are minimally upregulated in cells engineered with the triDrop.

### Functional CAR-T assays

A final series of experiments was performed to highlight the triDrop’s ability to engineer cells with immunotherapeutic capabilities. As a model system, T-cells were engineering to express an anti-CD19 CAR molecule (**Figure 5a**). Cells were electroporated using either manufacturer recommended conditions or 0.5 x 10^5^ cells per reaction with an anti-CD19 CAR mRNA payload.

**Figure 5.**
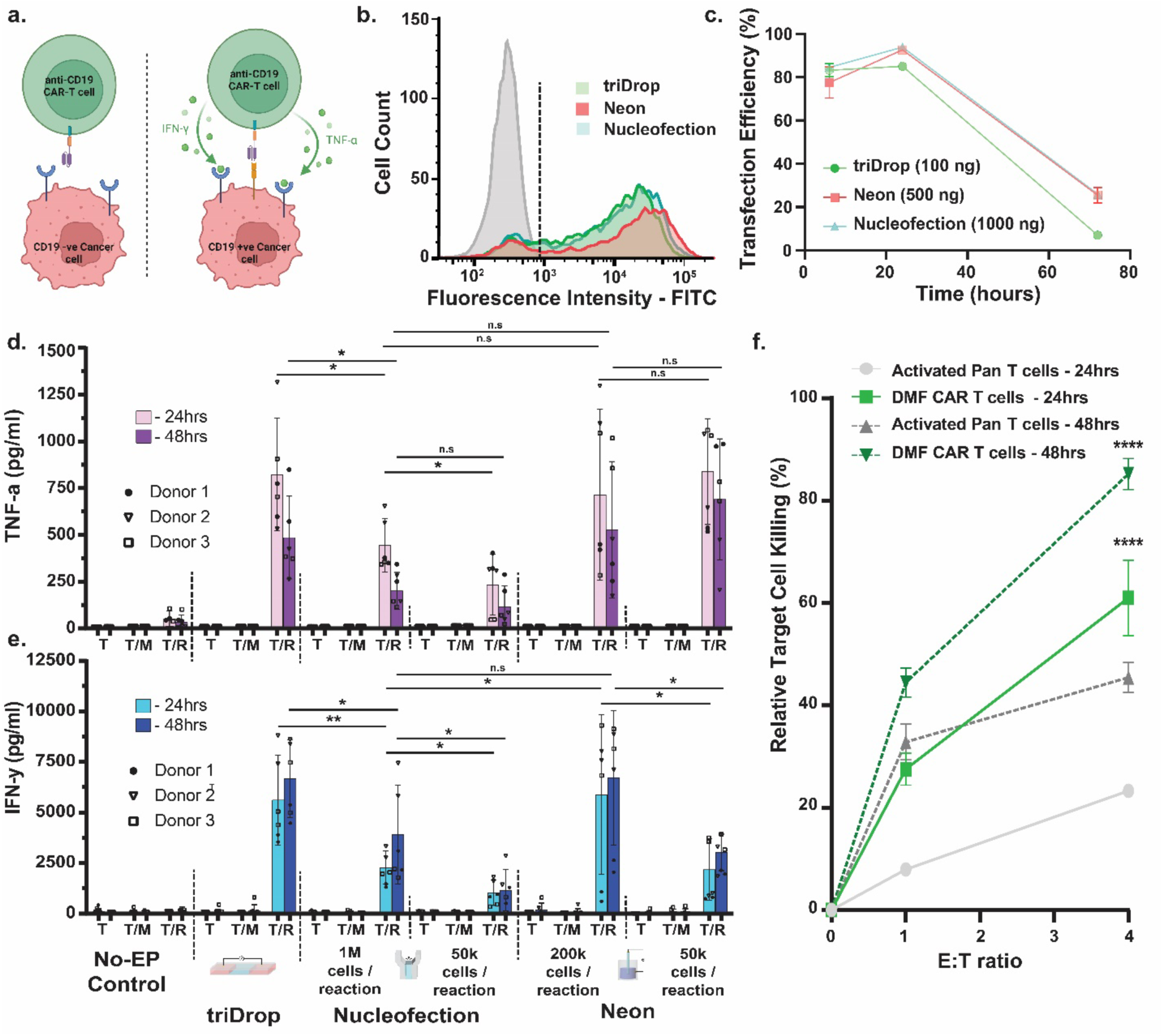
Functional CAR-T assays. a) Overview of immunotherapeutic testing assay detailing an anti-CD19 CAR T cell interacting with a CD19 negative or positive tumor cell line. Created with BioRender.com. b) Fluorescence intensity histograms showing FITC expressions for cells stained with a FITC-tagged CD19 protein from a control (grey), triDrop (green), Nucleofection (blue), and Neon (red). c) Line graphs depicting expression of an anti-CD19 CAR molecule at 6, 24, and 72 hours post EP when using the triDrop (green, 100ng of mRNA), Neon (red, 500ng of mRNA), and Nucleofection (blue, 1000ng of mRNA). Bar graphs depicting measured levels of d) TNF-α and e) IFN-γ after 24 hours (light purple and light blue) and 48 hours (dark purple and dark blue) of culture for cells electroporated using all three systems using either manufacturer recommended conditions or 0.5 x 10^5^ cells per reaction. Engineered cells are either cultured by themselves (T), or at a 1:1 ratio with MCF-7 cells (T/M), or with Raji cells (T/R). (n = 6) f) Line graphs depicting relative killing of Raji cells cocultured with activated Pan T cells at 1:1 ratio and 4:1 ratio after 24 hours (light grey, solid line) and 48 hours (dark grey, dashed line) or with triDrop engineered CAR T cells at a 1:1 and 4:1 ratio after 24 hours (light green, solid line), and 48 hours (dark green, dashed line). Statistical significance markers indicate difference between engineered cells and activated pan T cells for each timepoint. (n = 4). All error bars represent mean +/- SD. n.s indicated no significant difference, *, **,***, and **** represent p-values below 0.05, 0.01, 0.001 and 0.0001 respectively. Statistical analysis was performed using a student’s t-test.

We used optimal amounts of payload for each system (see **Figure 1**) and used flow cytometry to validate the delivery of the mRNA payload. **Figure 5b** shows the expression of the anti-CD19 protein by staining electroporated cells with a FITC-tagged CD19 protein. The expression of the CAR protein was > 80 % for all systems 6 hours post EP and peaked at 24 hours, then slowly declined over the subsequent two days (**Figure 5c**). Therefore, for the functionality assays, 6 hours post EP, 2.5 x 10^4^ engineered T-cells were cultured by themselves, or co-cultured with either 2.5 x 10^4^ MCF-7 cells (CD19-negative) or 2.5 x 10^4^ Raji cells (CD19-positive). **Figure 5 d and e** show bar graphs depicting the cytokine levels of IFN-γ and TNF-α in the supernatant for cells under various treatments and culture conditions after 24 hours and 48 hours of co-culture. None of the engineered cells produced either of the anti-tumour cytokines when cultured by themselves or in the presence of CD19 negative cells, indicating a highly specific response of the anti-CD19 CAR. When cultured with CD19-positive targets, cells engineered with the triDrop show significantly increased production of both IFN-γ (24 h: p = 0.006, 48 h: p = 0.05), and TNF-α (24 h: p = 0.02, 48 h: p = 0.016), compared to cells engineered with the Nucleofector when using with manufacturer recommend conditions. Cells engineered with the Neon also have higher cytokine production than Nucleofection but lack statistical significance across the 24 and 48 h timepoints. Additionally, using 0.5 x 10^5^ cells per reaction with the Nucleofector leads to a significant reduction in cytokine production capabilities. This aspect suggests that the Nucleofector impairs immune function immediately following EP (as seen by the transcriptional sequencing data) and that attempting to use fewer cells than recommended by the manufacturer further exacerbates this impairment. While the Neon was able to produce similar average levels of cytokines compared to triDrop, the response was highly variable across donors. Further, similar to the Nucleofector, using 0.5 x 10^5^ cells per reaction with the Neon surprisingly led to a significant reduction in the cells’ ability to produce IFN-γ (although not TNF-α). This suggests that, as hypothesized, using either the Neon or the Nucleofector with fewer than recommended cells lead to those cells having impaired functionality.

Finally, we test the ability to use cells engineered with our triDrop platform for killing Raji cells in co-culture. For this experiment, 2.5 x 10^4^ Raji cells were co-cultured with either 2.5 x 10^4^ or 10 x 10^4^ triDrop generated CAR T-cells or non-electroporated activated pan T cells. Coculture began 6 hours post EP. After 24 and 48 hours of co-culture, cells were collected and analyzed via flow cytometry by staining CD3 antibodies to separate target cells and effector cells and DAPI was used to differentiate living and dead cells. **Figure 5f** shows, as expected, the activated pan T cells elicit a small cytotoxic effect on the Raji cells, leading to ∼23 % (24h) and 45 % (48h) killing of target cells when co-cultured at the 4:1 effector: target ratio. However, when cultured with the triDrop engineered CAR T cells, approximately 27 % (24h) and 45 % (48h) of target cells are killed when cultured at a 1:1 ratio and 61 % (24h) and 85% (48h) are killed at a 4:1 effector: target ratio indicating a robust killing effect. Overall, the results detailed in **Figure 5** show that the triDrop platform can engineer cells capable of improved functional ability in immunotherapeutic assays while using less payload and fewer cells than either the Neon or Nucleofector EP systems. This capability will pave the way for high throughput immune cell engineering assays that can be both faster and more affordable than previous assays while producing reliable results.

### Conclusion

In this work we demonstrated that our miniaturized droplet electroporation system allows for a substantial reduction in the consumption of expensive reagents and precious cells compared to two state-of-the art electroporation system. We propose that this advancement will allow for a reduction in the cost of gene editing making cell therapy R&D more affordable, or alternatively making it possible to screen large libraries of edits on cells from rare cells or cells from a single donor. Additionally, we showed that the triDrop structure achieved on our DMF platform allows us to protect cells from the harmful effects of joule heating and electrochemical reactions that occur during the electroporation process and that this results in cells with less transcriptional dysregulation and improved functionality when compared to cells electroporated using traditional methods.

## Materials and Methods

DMF device fabrication and assembly, droplet operations, and the electroporation circuit (**Figure S3)** are described in the online supplementary information.

### Reagents and Materials

Unless specified otherwise, general-use chemicals and kits were purchased from Sigma-Aldrich (St. Louis, MO). Device fabrication reagents and supplies included chromium-coated glass slides, and gold-coated glass slides with AZ1500 photoresist from Telic (Valencia, CA), MF-321 positive photoresist developer from Rohm and Haas (Marlborough, MA), chromium etchant 9051 and gold etchant TFA from Transene (Danvers, MA), AZ-300T photoresist stripper from AZ Electronic Materials (Somerville, NJ), Teflon-AF 1600 from DuPont Fluoroproducts (Wilmington, DE). Transparency masks for device fabrication were printed from ARTNET Pro (San Jose, CA) and polylactic acid (PLA) material for 3D printing were purchased from 3Dshop (Mississauga, ON, Canada). General chemicals for tissue culture were purchased from Wisent Bio Products (Saint-Bruno, QC, Canada). Electronic components were obtained from DigiKey (Thief River Falls, MI). eGFP mRNA was purchased from TriLink Biotechnologies (catalog #: L-7201). Anti-CD19 CAR mRNA was purchased from Med Chem Express (catalog #: HY-153084). Cas9 and sgRNAs were purchased from Synthego (Redwood City, USA).

### Primary T-cell culturing

All primary T-cells were obtained from participants who gave informed consent, and had blood drawn in compliance with relevant ethical regulations under the approved summary protocol form 30009292 from the Ethics Committee at Concordia University. Primary human T cells were separated from fresh venous primary blood using LympPrep gradient centrifugation (Wisent Bio) and purified using either the EasySep Human CD4 T cell Isolation kit or EasySep Human T cell isolation kit to a purity of 95 % (STEMCELL Technologies, Canada, Catalog # 17952, and Catalog # 17951). All cells were kept in liquid nitrogen prior to use. Cells were thawed and cultured in complete culture medium consisting of RPMI-1640 with 10 % Fetal Bovine Serum (FBS), 100 IU / mL penicillin/streptomycin, and 200 IU / mL recombinant human IL-2 (STEMCELL Technologies, Canada, Catalog # 78220). Unless specified otherwise, cell culture was performed in U-bottom 96 well plates. After 24 hours post-thaw, the cells were activated with Human T-Activator CD3/CD28 Dynabeads (Fisher Scientific Ottawa, ON, #11131D) at a 1:1 cell to bead ratio and were incubated for 48 hours. After incubation, activator beads were removed following manufacturers protocol by first gently pipetting up and down to release cells from the activator beads followed by transferring the cells to a magnetic tube rack for 1-2 minutes to allow for cells and beads separation and the supernatant containing cells was transferred to a fresh tube. Cells were electroporated within 24 hours of bead removal. Primary T cells were counted using the ViCell Blu automated cell counter (Beckman Coulter, Canada) and maintained at 0.25 - 1 x 10^6^ cells / mL by daily addition of complete culture media. Post electroporation cells were recovered in the same media cocktail as above but with IL-2 levels increased to 400 IU / mL.

### Immortalized cell culture

Raji and MCF-7 cells were grown in cell culture media formed from RPMI (Raji) or DMEM (MCF-7), supplemented with 10% fetal bovine serum (FBS). Cells were grown to near confluency in complete media in T-25 flasks in an incubator at 37 °C with 5% CO2. Prior to each experiment, cells were detached using a solution of trypsin (0.25% w/v) and EDTA (1 mM), centrifuged, then resuspended in complete media. Cell lines were passaged every 2-3 days by media removal, PBS wash, trypsinization (for adherent cells), resuspension in fresh culture media and split at a 1:10 ratio into fresh media. All immortalized cell lines were kept below passage 10.

### Electroporation

For all electroporation platforms primary human T cells were electroporated within 24 hours after the removal of activation beads. Cells were in electroporation buffer for a maximum of 15 minutes and were transferred to recovery buffer promptly after pulse application using one of the three systems. For both the Neon and Nucleofector a wide range of pulse parameters (voltage amplitude, number of pulses, and pulse duration) have been shown for effective electroporation. In this work we use the pulse parameters recommended by the manufacturer for working with activated primary human T cells^65,66^ and shown to be effective by Zhang et al.^67^ and Schumann et al.^68^ which are three, 1600 VDC pulses, 10 ms in duration for the Neon and pulse code EO-115 for the Nucleofector.

### triDrop Electroporation

Detailed explanations of the triDrop platform can be found in our previous manuscript.^39^ Unless specified otherwise, 5 x 10^5^ cells were centrifuged (300 g, 5 min), washed in PBS, centrifuged, and resuspended in 10 µL of electroporation buffer with conductivity σ ∼ 8 mS / cm (Neon Type T buffer containing 0.05% Pluronic F68 surfactant) and payload was added as described below. 7.5 µL of a buffer with conductivity σ ∼ 16 mS / cm (PBS containing 0.05% Pluronic F68 surfactant) was pipetted into each of the device’s flanking reservoirs and 7.5 µL of the cell solution was pipetted into the middle reservoir. DMF actuation was used to dispense three unit droplets, ∼ 1 µL in volume, from each reservoir and actuate them to the three on-chip electroporation sites. Driving potentials (i.e. actuation) was used to merge the droplets into the triDrop configuration, and a pulse generation circuit was automatically triggered to deliver five, 550 VDC pulses, 500 µs in duration to the droplet structure. After electroporation cells were immediately removed from the chip via pipetting and transferred to recovery buffer. Cells from each of the three electroporation sites were loaded into their own well and served as technical triplicates.

### Neon Electroporation

Neon electroporation was performed using manufacturer recommended protocols for the Neon Transfection System 10 µL kit (Thermo Fisher, Catalog # MPK 1025). For each unique condition, 4.4 x 10^5^ cells were centrifuged (300 g, 5 min), washed in PBS, centrifuged again, and finally resuspended in 22 µL of type R buffer. 10 µL Neon tips were used a maximum of two times to serve as technical replicates. Three, 1600 VDC pulses, 10 ms in duration were applied. Immediately, electroporation cells were transferred to recovery buffer.

To explore the effects of cell density on electroporation performance, 8 x 10^5^ cells were washed in PBS and resuspended in 40 µL of Buffer R. The cell solution was diluted with fresh Buffer R in the following ratios, 5.5 µL: 16.5 µL and 11 µL: 11 µL, creating aliquots containing 1.1 x 10^5^ cells and 2.2 x 10^5^ cells respectively. These aliquots were used to perform two electroporation reactions each containing ∼ 5 x 10^4^ cells per reactions and ∼ 1 x 10^5^ cells per reaction respectively. The remaining cell solution was used to perform two reactions with 2 x 10^5^ cells per reaction (manufacturer recommended amount). All conditions were cultured at 5 x 10^4^ cells per well for post-electroporation analysis.

### Lonza Electroporation

Lonza electroporation was performed using manufacturer recommended protocols for the P3 Primary Cell 4D-Nucleofector X Kit S (Lonza, Canada, Catalog # V4XP-3032). For each condition, 1.1 x 10^6^ cells were centrifuged (300 g, 5 min), washed in PBS, centrifuged, and resuspended in 22 µL of P3 buffer. 20 µL of cell solution was pipetted into the nucleofector cuvette and pulse code E0-115 was used for electroporation. Immediately following electroporation 100 µL of pre-warmed recovery buffer was added to each cuvette and the sample was incubated for 10 minutes prior to be transferred to a well plate.

To determine the effects of cell density on electroporation efficacy, 3 x 10^6^ cells were washed in PBS and resuspended in 60 µL of P3 buffer. The cell solution was mixed with fresh P3 buffer in the following ratios, 2:38, 4:36, and 8:32, creating aliquots containing 1 x 10^5^ cells, 2 x 10^5^ cells, and 4 x 10^5^ cells. These aliquots were each used to perform two electroporation reactions containing 5 x 10^4^ cells per reaction, 1 x 10^5^ cells per reaction or 2 x 10^5^ cells per reactions respectively. The remaining cell solution was used to perform two reactions with 1 x 10^6^ cells per reaction (manufacturer recommended amount). All conditions were cultured at 50,000 cells per well for post-electroporation analysis.

### Transfection protocols

#### mRNA transfection

mRNA transfection was performed by preparing cells for electroporation using the techniques outlined above and adding mRNA immediately prior to electroporation. Unless specified otherwise, mRNA was added at 50 ng / µL for each system (50 ng per reaction for triDrop, 500 ng per reaction for Neon, and 1000 ng per reaction for Lonza). eGFP and anti-CD19 CAR mRNA were stored at −80 ^°^C in 5 µL aliquots with a stock concentration of 1 and 2 µg / µL respectively. The eGFP-mRNA sequence can be found in **Table S2**.

For experiments exploring the effects of mRNA concentration on electroporation efficacy, 100 ng, 50 ng, and 10 ng per reaction conditions were explored using the triDrop. Three tubes were prepared, each containing 5 x 10^5^ cells resuspended in 10 µL of electroporation buffer as described above. 1 µL of stock eGFP-mRNA (1 µg / µL) was added to the first tube, 0.5 µL of stock solution was added to the second tube, and for the final tube 1 µL of stock mRNA was diluted with 9 µL of type T electroporation buffer and 1 µL of the dilution was added to the cell mixture. For Neon, 500 ng, 100 ng, and 50 ng reaction conditions were explored. Three tubes each containing 4.4 x 10^5^ cells resuspended in 22 µL of electroporation buffer were prepared as outlined above. 1 µL of stock mRNA (1 µg / µL) was added to the first tube. Next, 1 µL of stock mRNA was diluted in 9 µL of type R electroporation buffer. 2 µL of this dilution was added to the second tube, and 1 µL was added to the third tube. Each tube was used for two electroporation reactions.

For the Lonza, 500 ng, 100 ng, and 50 ng per reaction conditions were explored. Three tubes each containing 2.2 x 10^6^ cells resuspended in 44 µL of electroporation buffer was prepared as outlined above. 1 µL of stock mRNA (1 µg / µL) was added to the first tube. Next, 1 µL of stock mRNA was diluted in 9 µL of P3 electroporation buffer. 2 µL of this dilution was added to the second tube, and 1 µL was added to the third tube. Each aliquot was used for two electroporation reactions.

#### Gene Editing

TRAC knockouts were performed by complexing TrueCut Cas9 protein V2 (Thermo Fisher, Canada, catalog #: A36496) with an sgRNA cocktail designed by Synthego to target the TRAC locus (Synthego, SKU: 052-1020-000-1.5n-0). Cas9 proteins and gRNAs were kept at −20 °C at a stock concentration of 30 pmol / µL and 100 pmol / µL respectively. The amount of gRNA and Cas9 protein added to each sample was controlled by either normalizing to the number of cells per reaction or by normalizing to the volume of the reaction. For normalization to the number of cells per reaction, 50 pmol of Cas9 was mixed with 100 pmol of gRNA for every 1 million cells used. For normalization to reaction volume, 50 pmol of Cas9 was mixed with 100 pmol of gRNA for every 20 µL of reaction volume (equivalent to volume required for one Lonza nucleofection cuvette). After mixing of the Cas9 protein with the gRNA, the mixture was left at room temperature for 10 minutes to allow for the formation of the ribonucleic protein (RNP) complex and either immediately added to the cell mixture or kept on ice for no more than 30 minutes before being added to the cell mixture.

CRISPR knock-ins were performed by using the protocol originally described by Cloarec-Ung et al..^51^ crRNA (200 pmol / µL, IDT) was mixed with tracrRNA (200 pmol / µL, IDT) at a 1:1 ratio and annealed by heating at 95 °C for 5 minutes followed by cooling to room temperature at a rate of 0.1 °C / s in a thermocycler to form sgRNA at a concentration of 100 pmol / µL. the resulting sgRNA was then mixed with Cas9 (30 pmol / µL) at a 2:1 ratio and left to complex for 10 minutes at room temperature to form RNPs. A single stranded oligonucleotide (ssODN) HDR repair template (400 pmol / µL, IDT) was diluted 1:8 in electroporation buffer prior to electroporation. p53 siRNA (100 pmol / µL, Thermo Fisher, Catalog # 4390824) was diluted 1:1000 in nuclease free water. For every 20 µL of reaction volume, 50 pmol of RNP, 50 pmol of ssODN, and 20 fmol of p53 siRNA was added to cells already in electroporation buffer immediately prior to electroporation. Following electroporation cells were recovered in the recovery buffer described above with the addition of 0.5 µM AZD7648 (Cayman Chemical, Item # 28598).

### Genetic Analysis

#### RT-qPCR

RT-qPCR was performed in accordance with the Minimum Information for Publication of Quantitative Real Team PCR Experiments (MIQE) guidelines^69^ and associated data (including primer sequences, qPCR validation, and data analysis) can be found in **Figure S4** and **Table S5**. RT-qPCR was performed on the Eco Real Time PCR System (catalog # EC-101-1001) using the Eco study V5.0 software. Cells from two donors were electroporated in technical replicates with each system using the methods outlined above using either the manufacturer recommended conditions or 5 x 10^4^ cells per reaction. Post electroporation all cells were cultured at 5 x 10^4^ cells cells per well in 200 µL of recovery buffer (recipe above). 6 hours after electroporation cells were washed in PBS, lysed and prepared for RT-qPCR using Power SYBR green cells-to-Ct kit (Thermo Fisher, Catalog #4402954). PCR cycling conditions were as follows: 95 °C, 10 min; 40x (95 C° 15s, 60 C° 1 min). 18s rRNA was used as a reference gene.^70,71^

#### Knock-In

48-hours post-electroporation genomic DNA (gDNA) was recovered from the cells using the DNeasy Blood and Tissue Kit (Qiagen, Catalog # 69504). gDNA was amplified using 0.5 µM of pri077F+R primers (**Table S2**) with the following thermal cycle: 98 C° 30s; 35x (98 C° 10s, 60 C° 10s, 72 C° 30s); 72 C° 5 min. Following amplification, PCR products were purified with the GeneJET PCR purification kit (Thermo Fisher, Catalog #K0702). Purified PCR products were quantified by Nanodrop at a range of 20 – 40 ng / µL, and 5-15 ng were sent for Sanger sequencing at the IRIC Genomics core with the primer pri0003-A1(**Table S2**). Chromatograms for each condition were aligned against a control sample using SnapGene’s alignment tool and validated with Synthego’s ICE Analysis. (**Figure S5**).

#### Transcriptomic sequencing and analysis

Transcriptomic sequencing was performed using the Oxford Nanopore minION Mk1B with flow cell R9.4.1. Cells from three donors were electroporated using the optimized or manufacturer (Neon and Nucleofector) recommended condition. All cells post-electroporation were cultured at 5 x 10^4^ cells per well in 200 µL of recovery buffer. Six hours post electroporation cells were washed in PBS, lysed, and RNA was extracted using RNeasy Mini Kit (Qiagen, catalog #74104). As per manufacturer’s recommendations, 200 ng of total RNA was used from each condition as an input to the kit with an RNA integrity number > 10. The RNA library was prepared using the PCR-cDNA Barcoding Kit (Oxford Nanopore, catalog #SQK-PCB111.24) following the manufacturer’s protocol. Briefly, by taking full-length polyadenylated RNA, complementary strand synthesis and strand switching were performed using kit-supplied oligonucleotides. The kit contained 24 primer pairs, 12 of which were used to generate and then amplify double-stranded cDNA by PCR amplification using rapid attachment barcode primers with the following thermal cycle: 95 C° 30s; 18x (95 C° 15s, 62 C° 15s, 65 C° 120s), 65 C° 360s, 4 C° hold. These primers contained 5’ tags and facilitated the ligase-free attachment of Rapid Sequencing Adapters and their sequences are listed in the **Table S2**. 1 µL of amplified cDNA from each condition was analyzed using a 4200 Tapestation System (Agilent) prior to sequencing to confirm sample quality and a concentration of at least 1000 pg / µL. Amplified and barcoded samples were then pooled together, and Rapid Sequencing Adapters were added to the pooled mix. The final pooled library loaded on to the flow cell contained 2 fmol of cDNA from each barcoded condition for a total of 24 fmol.

Base calling was performed using Oxford Nanopore’s Dorado software. Sequencing reads were aligned with the Homo Sapiens GRCh38 transcriptome using Minimap2.^72^ Transcript counts were performed using Salmon^73^ and mapped to genomic data using the R/Bioconductor package biomaRt.^74^ Differential expression was calculated using DESeq2.^75^ Z – scores were used to quantify variation from the control based on the number of transcripts per million (TPM) that were counted for each gene using the following formula:

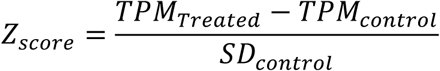

Genes were grouped into high level genetic pathways using the Reactome data base^62^ and analyzed using the ReactomePA package.^76^

### Flow Cytometry

Flow cytometry was performed using the BD FACS Melody (BD Bioscience, Canada). The FACS was equipped with three excitation lasers (405 nm, 488 nm, 561 nm) in a 2B-2V-4YG configuration. TRAC knockouts were detected via staining with anti-human alpha/beta TCR FITC (Thermo Fisher Scientific, catalog #11-9955-42). Anti-CD19 CAR expression was detected using FITC-labelled human CD19 protein (Acro Biosystems, catalog # CD9-HF251-25ug). CD3 expression was detected using PE-CD3 monoclonal antibody (Thermo Fisher Scientific, catalog #12-0038-42). CD19 expression was detected using a FITC Mouse anti-Human CD19 antibody (BD Bioscience, catalog #560994). All staining was performed in accordance with manufacturer guidelines, cells were incubated with the appropriate stains for 30 – 60 minutes at 4 ^°^C followed by three PBS wash steps to remove any unbound antibodies. For all samples, viability was assessed by staining dead cells using DAPI (50 ng / µL) added to the sample immediately prior to FACS and mixed thoroughly with the sample by pipetting. Lymphocytes were separated from debris using forward scatter vs side scatter dot plots and singlets were separated from doublets by plotting side scatter – pulse width vs side scatter – pulse height.

### CAR-T assays

mRNA coding for a second-generation anti-CD19 CAR molecule with a 41BB-CD3z coactivation domain was used. For each electroporation system, > 4 x 10^5^ activated pan T cells were electroporated with anti-CD19 CAR mRNA using either the manufacturer recommended conditions or by performing eight reactions using 0.5 x 10^5^ cells each (1 x 10^6^ cells were used for the Nucelofector). Immediately post electroporation cells were incubated in 200 µL recovery buffer at 0.5 x 10^5^ cells per well.

Four hours prior to the tumour cell killing assay, 2.5 x 10^4^ Raji cells per well were seeded in a U-bottom well plate in 100 µL of fresh RPMI with 10 % FBS and 2.5 x 10^4^ MCF-7 cells per well were seeded in a treated flat-bottom 96 well plate in 100 µL of fresh RPMI with 10 % FBS. Control wells were seeded with 100 µL of fresh RPMI containing no cells.

After six hours of recovery, electroporated and control cells were counted, washed in PBS, and resuspended in fresh RPMI with 10% FBS at a concentration of 2.5 x 10^4^ cells per 100 µL. 100 µL of cell solution form each condition was then added in technical replicates to wells prepared earlier containing either just media, MCF-7 cells, or Raji cells. Supernatant was collected after 24-hours and 48-hours by centrifuging the plates for 3 min at 300 g to pellet the cells and pipetting 180 µL from the top of each well. Supernatant was analyzed via ELISA using the manufacturer recommended protocol for INF-γ (BD Biosciences, catalog # 555142) and TNF-α (BD Biosciences, catalog # 555212).

Tumour cell killing was performed by co-culturing CAR T cells or control activated pan T cells with Raji cells at a 1:1 (2.5 x 10^4^ T cells: 2.5 x 10^4^ Raji cells) or 4:1 ratio (1 x 10^5^ T cells: 2.5 x 10^4^ Raji cells). Raji cell death was validated via flow cytometry.^77^ After 24-hours of co-culture, cells are recovered and washed before being stained with PE-CD3 monoclonal antibody and prepared for flow cytometry. Immediately before flow cytometry DAPI is added to visualize cell viability. PE-CD3 staining allows for the differentiation between T cells and Raji cells, and DAPI allows for the visualization of living and dead cells. A detailed gating overview is shown in X **Figure S6**. Relative Killing Efficiency was measure using the following formula:

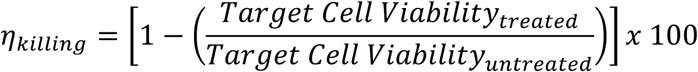

Where the treated target cell viability represents the viability of Raji cells cocultured with either activated pan T cells or CAR T cells, and the untreated viability represents the viability of Raji cell cultured by themselves under the same conditions over the same period.

## Acknowledgments

FMCU and DJHFK thank the Terry Fox Research Institute (Terry Fox New Investigator Award #TFRI 1118). DJHFK has salary support from FRQS in the form of a Chercheurs-boursiers Junior 1 fellowship (#283502). FMCU was supported by a Cole Foundation Doctoral Award, a bourse d’excellence du programme de biologie moléculaire from the Université de Montréal and a PhD scholarship from the Institut de Recherche en Immunologie et en Cancérologie. SCCS thank the Natural Sciences and Engineering Research Council (NSERC), the Fonds de Recherche Nature et Technologies (FRQNT), the Canadian Foundation of Innovation (CFI), and MEDTEQ for funding. SRL thanks NSERC CGS-D for funding. SRL and NR thank Concordia University Department of Electrical and Computer Engineering for FRS funding. SCCS thanks Concordia University for a Research Chair.

## Author Contributions

Experiments were designed by SRL, and SCCS. SRL and NR performed transcriptomic analysis. NR and JP designed qPCR primers, qPCR was optimized and performed by NR. DMF control software along with electroporation hardware and software were developed by SRL. FG, MH, and PJD developed the methodology for cell isolation and freezing from fresh blood and FG and MH performed the isolation and freezing protocols as well as all ELISAs. FMCU and DJHFK developed CRISPR knock in methodology. SCCS secured required funding for the project. SRL and SCCS wrote the manuscript which was revised and reviewed by all authors.

