## Supplementary material for "A Digital Microfluidic Platform for the Microscale Production of Functional Immune Cell Therapies": SI

### Supplementary Methods

##### DMF ‘tri-drop’ device fabrication

triDrop devices were composed of a top plate and a bottom plate. The bottom plate consisted of chromium electrodes divided into an array of 30 actuation electrodes (2 mm by 2 mm), 12 reservoir electrodes (2.9 mm by 5.5 mm) arranged into 3 reservoirs, 6 active dispensing electrodes (2 mm by 2 mm), and 3 splitting electrodes (3.8 mm x 3 mm). All electrodes were separated by a 150 µm gap. Bottom plates were coated in an SU8-5 dielectric layer (~ 5 µm thickness) and further coated in Teflon-AF 1600 in 2 % w/w in Fluorinert FC-40 to serve as a hydrophobic layer.

TriDrop top plates bearing gold electrodes (0.2 mm wide) were formed from a glass substrate coated with 100 nm gold adhered to seed chromium layer (~12 nm). To form the gold electrodes, top plates were spin-coated (10 s 500 rpm, 30 s 3000 rpm, 20 s 5000 rpm) in S1811, exposed through a transparent mask, developed using Microposit MF321 (2 min), washed with DI water, submerged in gold etchant (2 min), washed with DI water, and submerged in AZ stripper to remove the remaining photoresist before being washed with acetone, IPA, and DI water, and dried with nitrogen. To disconnect the chromium from the gold wiring, we followed the above protocol except using CR-4 etchant to remove the chromium. To insulate the gold electroporation electrodes from the Cr-grounding layer, the top plate was surface treated for 45 s in a plasma cleaner (Harrick Plasma PDC-001, Ithaca, NY) before coating a 5 µm dielectric of SU8-5. Briefly, the photoresist was spin-coated (10 s 500 rpm, 30 s 2500 rpm), followed by a soft bake (65 °C 2 min, 95 °C, 10 min), exposed to UV light through a custom mask (5 s), post-exposure baked (65 °C 2 min, 95 °C 10 min), developed in SU8 developer (15 s), rinsed with IPA and DI water, dried with nitrogen, and then hard baked (180 °C, 10 min). Top plates were spin-coated with Teflon-AF

1600 in 2 % w/w in Fluorinert FC-40 (10 s 500rpm, 30s 1500rpm). To assemble the completed triDrop device, the top and bottom plates were assembled using two layers of double-sided tape (180 μm total thickness, 3M) and the gold electrode on the top plate were aligned directly above electroporation sites on the bottom plate.

**DMF 'tri-drop' device assembly**

The bottom plate of the triDrop device was placed on a pogo pin holder that has been propped to a height 20 cm above the benchtop using a chassis constructed from T-slotted aluminum extrusions purchased from McMaster-Carr (catalog #: 47065T101, Aurora, OH) and machined and assembled in-house. The system is connected to a 720 pixel, 30 frames-per-second camera (Skybasic, Houston TX.) to visualize droplet movements on the device. A 12-input card edge connector from Digikey (catalog #: 151-1410-ND, Thief River Falls, MI), was attached to the top plate of the triDrop device and connected via three leads (DMF ground, High Voltage DC, DC ground). The two DC leads were connected to an electroporation pulse circuit and one lead was used to provide the electrical connection for the DMF ground. The electroporation circuit consisted of an 8 pin optocoupler (Model #: AQW216EH) purchased from Digikey was connected to a Z650-0.32-U DC power source (TDK-Lambda) and controlled by an Arduino Uno running a custom pulse generating program, creating custom pulses of varying amplitudes and durations (100 - 600 $V_{DC}$, 0.2-10 ms in duration).

**DMF 'tri-drop' device operation**

For automating droplet movement on the device, see our previous published work for circuit and connectivity.[1] The electroporation and DMF actuation circuit were controlled by our in-house software which is available on our bitbucket registry (https://bitbucket.org/shihmicrolab/littleleung_2023). Droplet movements were programmed by application of AC potentials (300 – 400 $V_{RMS}$) at 15 kHz between the top and bottom plates. The DMF actuation software was also used to initiate the electroporation pulse circuitry (**Figure S3**) to ensure immediate and uniform pulse application after triDrop merging.

#### Supplementary Tables

**Table S1: Cost of Genetically Engineering Primary Human T cells**

| Process | Material | Material Cost (CAD) | Nucleofector | | Neon | | triDrop | |
|---|---|---|---|---|---|---|---|---|
| | | | Amount Needed per reaction | Cost (CAD) | Amount Needed per reaction | Cost (CAD) | Amount Needed per reaction | Cost (CAD) |
| **CD4+ T Cell Isolation** | Leukopack | 4757 (~ 5 x $10^9$ cells) | 1 x $10^6$ cells | 0.95 | 2 x $10^5$ cells | 0.19 | 5 x $10^4$ cells | 0.05 |
| | Isolation Kit | 979 (1 x $10^9$ cells) | 1 x $10^6$ cells | 0.98 | 2 x $10^5$ cells | 0.2 | 5 x $10^4$ cells | 0.05 |
| **Cell Culture** | Media + Serum w/ cytokines | 85 (500 mL) | 1 mL | 0.17 | 0.2 mL | 0.03 | 0.05 mL | <0.01 |
| | Activation Beads | 1195 (2 mL) | 0.025 mL | 15 | 0.005 mL | 3 | 0.00125 mL | 0.75 |
| **Gene Editing** | Cas9 Nuclease | 1294 (500 µg/3nmol) | 50 pmol | 21.50 | 10 pmol | 4.32 | 2.5 pmol | 1.08 |
| | sgRNA | 95 (1.5 nmol) | 100 pmol | 6.33 | 20 pmol | 1.24 | 5 pmol | 0.31 |
| **TOTAL COST FOR 1 REACTION** | | | **$44.90** | | **$8.98** | | **$2.25** | |

**NOTE**: Reagent costs are validated as of June 2024 and may change with bulk purchasing or institutional pricing. It is assumed that ~50% of cells in a Leukopak are pan T cells. Costs will change depending on what cell line is being engineered. Leukopak – StemCell Technologies (catalog # 70500); Isolation Kit – StemCell Technologies (catalog # 17951); serum – Thermo Fisher (catalog # A5670701); cytokines – Thermo Fisher (catalog #200-02-1MG); activation beads – Thermo Fisher (catalog #11161D); Cas9 Nuclease - Thermo Fisher (catalog #A36496); gRNA – Synthego (SKU: SKU: 052-1020-000-1.5n-0)

**Table S2: Important Genetic Sequences**

| Name | Sequence |
| --- | --- |
| eGFP mRNA | AUGGUGAGCAAGGGCGAGGAGCUGUUCACCGGGGUGGUGCCCAUCCUGGUC GAGCUGGACGGCGACGUAAACGGCCACAAGUUCAGCGUGUCCGGCGAGGGC GAGGGCGAUGCCACCUACGGCAAGCUGACCCUGAAGUUCAUCUGCACCACCG GCAAGCUGCCCGUGCCCUGGCCCACCCUCGUGACCACCCUGACCUACGGCGU GCAGUGCUUCAGCCGCUACCCCGACCACAUGAAGCAGCACGACUUCUUCAAG UCCGCCAUGCCCGAAGGCUACGUCCAGGAGCGCACCAUCUUCUUCAAGGACG ACGGCAACUACAAGACCCGCGCCGAGGUGAAGUUCGAGGGCGACACCCUGG UGAACCGCAUCGAGCUGAAGGGCAUCGACUUCAAGGAGGACGGCAACAUCC UGGGGCACAAGCUGGAGUACAACUACAACAGCCACAACGUCUAUAUCAUGG CCGACAAGCAGAAGAACGGCAUCAAGGUGAACUUCAAGAUCCGCCACAACA UCGAGGACGGCAGCGUGCAGCUCGCCGACCACUACCAGCAGAACACCCCCAU CGGCGACGGCCCCGUGCUGCUGCCCGACAACCACUACCUGAGCACCCAGUCC GCCCUGAGCAAAGACCCCAACGAGAAGCGCGAUCACAUGGUCCUGCUGGAG UUCGUGACCGCCGCCGGGAUCACUCUCGGCAUGGACGAGCUGUACAAGUAA |
| SRSF2 gRNA | /AlTR1/rCrGrGrCrUrGrUrGrGrUrGrUrGrArGrUrCrCrGrGrGrUrUrUrArGrArGrCr UrArUrGrCrU/AlTR2/ |
| ssODN HDR template | T*G*GACGGCCGCGAGCTGCGGGTGCAAATGGCGCGCTACGGCCGCCCTCCAG ATTCACACCACAGCCGCCGGGGACCGCCACCCCGCAG*G*T |
| Pri077 F | AGCGATATAAACGGGCGCAG |
| Pri077 R | TCGCGACCTGGATTTGGATT |
| Pri0003-A1 | T*C*GGCGACGTGTACATCC |
| Barcode Flanking Sequence (Top Strand) | 5' - ATCGCCTACCGTGA - barcode - TTGCCTGTCGCTCTATCTTC - 3' |
| Barcode Flanking Sequence (Bottom Strand) | 5' - ATCGCCTACCGTGA - barcode - TCTGTTGGTGCTGATATTGC - 3' |
| Barcode 01 | AAGAAAGTTGTCGGTGTCTTTGTG |
| Barcode 02 | TCGATTCCGTTTGTAGTCGTCTGT |
| Barcode 03 | GAGTCTTGTGTCCCAGTTACCAGG |
| Barcode 04 | TTCGGATTCTATCGTGTTTCCCTA |
| Barcode 05 | CTTGTCCAGGGTTTGTGTAACCTT |
| Barcode 06 | TTCTCGCAAAGGCAGAAAGTAGTC |
| Barcode 07 | GTGTTACCGTGGGAATGAATCCTT |
| Barcode 08 | TTCAGGGAACAAACCAAGTTACGT |
| Barcode 09 | AACTAGGCACAGCGAGTCTTGGTT |
| Barcode 10 | AAGCGTTGAAACCTTTGTCCTCTC |
| Barcode 11 | GTTTCATCTATCGGAGGGAATGGA |
| Barcode 12 | CAGGTAGAAAGAAGCAGAATCGGA |

**Table S3: Mutually Dysregulated Genes ($p < 0.05$)**

| Gene Symbol | Gene Name | Ensembl Gene ID | Fold Change | | | Putative Function |
|---|---|---|---|---|---|---|
| | | | triDrop | Neon | Nucleofection | |
| ACBD3 | acyl-CoA binding domain containing 3 | ENSG00000182827 | 3.71228801 | 3.77293021 | 4.26957006 | Involved in the maintenance of Golgi structure and function |
| MT1M | Metallothionein 1M | ENSG00000205364 | 2.87867827 | 2.77319454 | 3.70987145 | Member of the metallothionein superfamily |
| SLC7A11 | Solute carrier family 7 member 11 | ENSG00000151012 | 4.36751455 | 4.49961256 | 3.7043796 | Implicated in ferroptosis[2] |
| PMCH | Pro-Melanin Concentrating Hormone | ENSG00000183395 | 1.59762742 | 2.38135295 | 2.6530848 | Expression in immune cells may inhibit proliferation[3] |
| CRIP1 | Cysteine rich protein 1 | ENSG00000213145 | 1.17007127 | 1.3629836 | 1.80771559 | LIM/double zinc finger protein family |
| RPL38P4 | Ribosomal Protein L38 Pseudogene 4 | ENSG00000250562 | 1.19873041 | 1.38231401 | 1.49196731 | Pseudogene |

**Table S4: Reactome Pathway Analysis**

| Reactome | | Z-scores | | |
|---|---|---|---|---|
| **Pathway ID** | **Brief description** | **Nucelofection** | **triDrop** | **Neon** |
| R-HSA-5661231 | Metallothioneins bind metals | 7.857664356 | 3.423042716 | 9.99969 |
| R-HSA-5660526 | Response to metal ions | 6.395220981 | 2.653043923 | 7.833048 |
| R-HSA-2028269 | regulation of cell proliferation and apoptosis | 4.41568806 | 1.176575666 | 4.359968 |
| R-HSA-9614657 | transcription of cell death genes | 2.637279679 | 0.401525923 | 2.575192 |
| R-HSA-9635465 | suppression of apoptosis | 2.544866109 | 1.800080837 | 2.624122 |
| R-HSA-111458 | formation of apoptosome | 2.210876005 | 1.340514592 | 2.419572 |
| R-HSA-9818027 | cytoprotective genes | 3.355447126 | 2.238439352 | 3.910091 |
| R-HSA-9818035 | endoplasmic reticulum stress associated genes | 3.99123084 | -0.123698357 | 2.488199 |
| R-HSA-9648895 | integrated stress response phosphorylation | 3.915343024 | 1.7811285 | 4.282006 |
| R-HSA-111995 | lipid mediators involved in inflammatory responses | 7.810607731 | 4.059175677 | 8.231199 |
| R-HSA-9818026 | inflammation associated genes | 3.683541285 | -0.287309368 | 2.024568 |
| R-HSA-75876 | resolution of inflammatory response | 1.643061815 | 0.390802551 | 4.152934 |

#### Table S5: qPCR Primer Sequences

| Gene Symbol | Gene Name | Ensemble ID | Primer sequence | PCR product (bp) | Product Melting Point (°C) |
|---|---|---|---|---|---|
| 18srRNA | 18s ribosomal RNA | NCBI ID: 106631781 | **F:** 5'-CTC AAC ACG GGA AAC CTCAC-3'<br>**R:** 5'-CGC TCC ACCAAC TAA GAACG-3' | 110 | 82.3 |
| IL-2 | Interleukin-2 | ENSG00000109 471 | **F:** 5'-CACAGCTACAACTGGAGCATTTAC-3'<br>**R:** 5'-TTCAGTTCTGTGGCCTTCTTGG-3' | 133 | 76.6 |
| IFNG | Interferon gamma | ENSG00000111 537 | **F:** 5'- GAGTGTGGAGACCATCAAGGA -3'<br>**R:** 5'-GGACATTCAAGTCAGTTACCGAA-3' | 113 | 78.1 |
| TNFα | Tumor necrosis factor | ENSG00000232 810 | **F:** 5'- AACCTCCTCTCTGCCATCAA -3'<br>**R:** 5'-GGAAGACCCCTCCCAGATAG -3' | 100 | 84 |

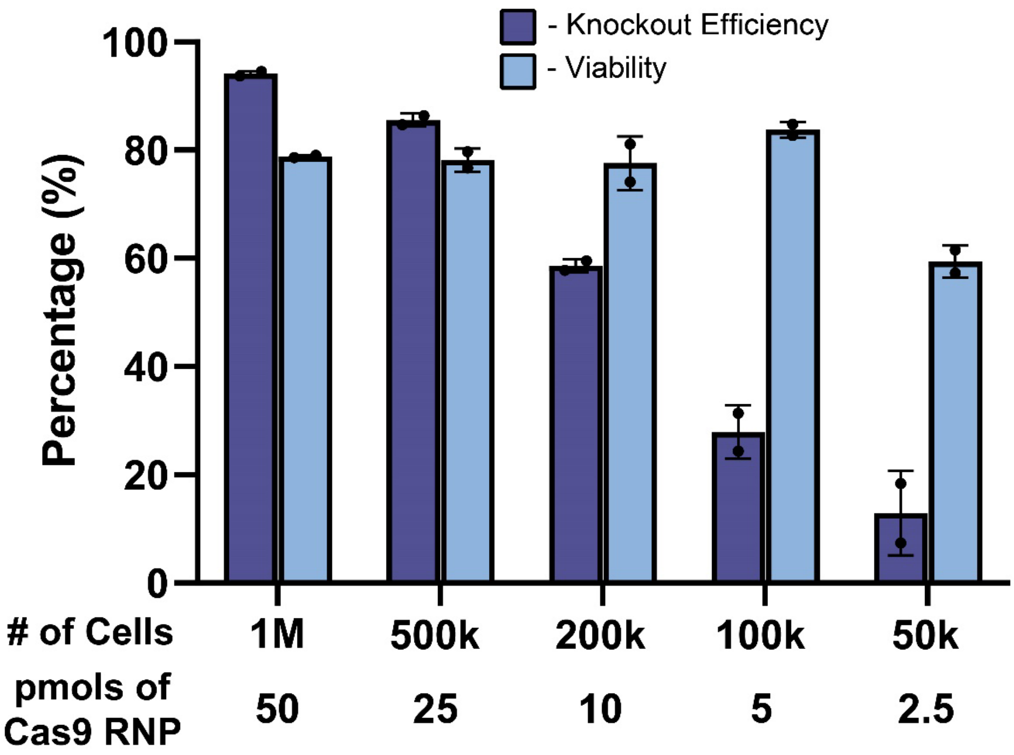


**Figure S1. Reducing Payload and Cell Count with the Nucleofector.** Bar graphs showing knockout efficiency (dark blue) and viability (light blue) measured four days post-EP for cells engineered with the Nucleofector. The number of cells and amount payload are indicated under each set of bars. All error bars represent mean +/- 1 SD.

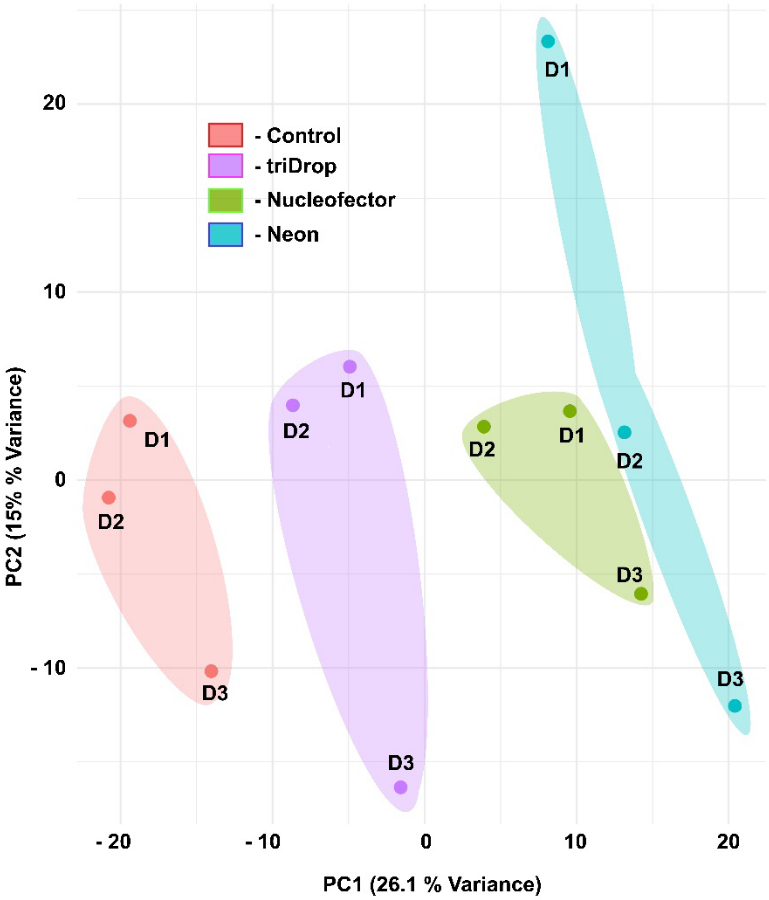


**Figure S2. RNAseq data.** Principal component analysis performed across all genes for three donors electroporated with the three EP systems. Control (red cluster), triDrop (purple cluster), Nucleofector (green cluster) and Neon (blue cluster) are all shown. Cluster proximity indicates increased similarity between samples meaning the triDrop is most similar to the control six hours post EP. Graph was generated using DESeq2.

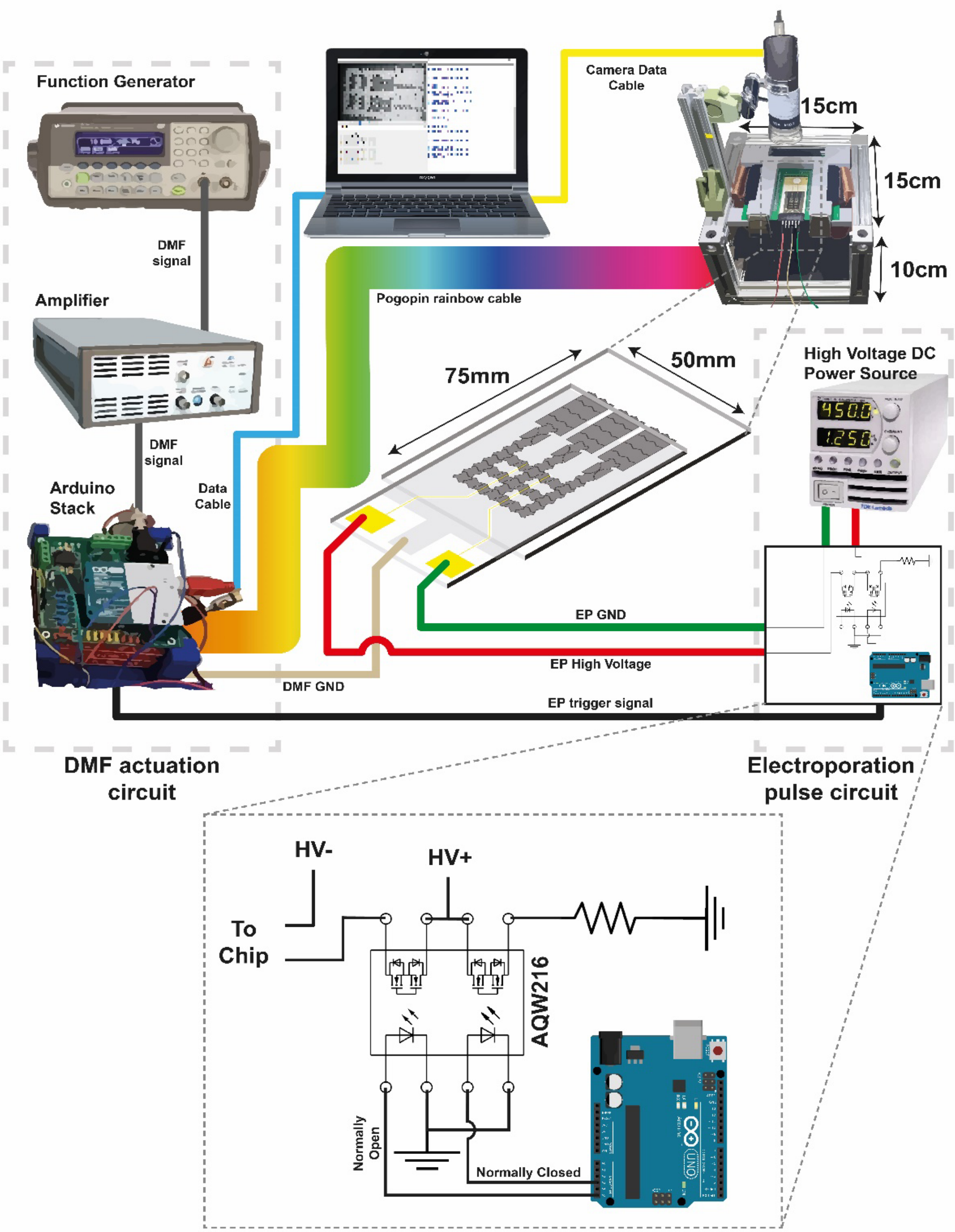


**Figure S3. DMF and electroporation circuit diagram.** Schematic overview of the complete triDrop automation setup detailing the DMF actuation hardware, automated electroporation pulse generation circuit, and chip holder.

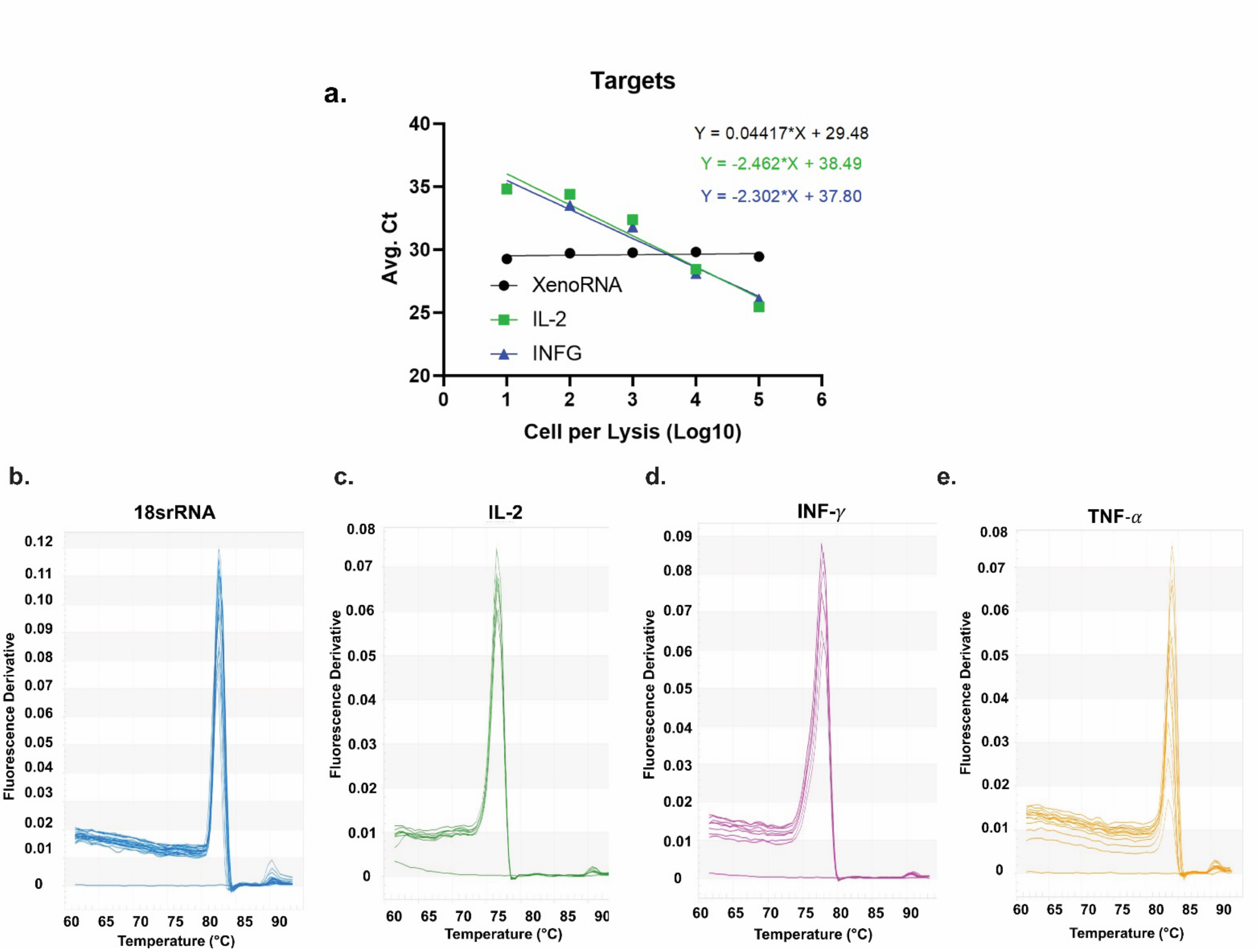


**Figure S4. qPCR raw data. a**) Standard curve depicting relationship between threshold cycle and number of lysed cells. Melting curves for b) 18srRNA, c) IL-2, d) IFN-γ, e) TNF-α showing product melt point consistent with that predicted in **Table S5.**

**Wild Type Sequence**

tggacggccgcgagctgcgggtgcaaatggcgcgctacggccgccccccggactcacaccacagccgccggggaccgccaccccgcaggt

**ssODN HDR Template Sequence**

tggacggccgcgagctgcgggtgcaaatggcgcgctacggccgccc**T**cc**A**ga**T**tcacaccacagccgccggggaccgccaccccgcaggt

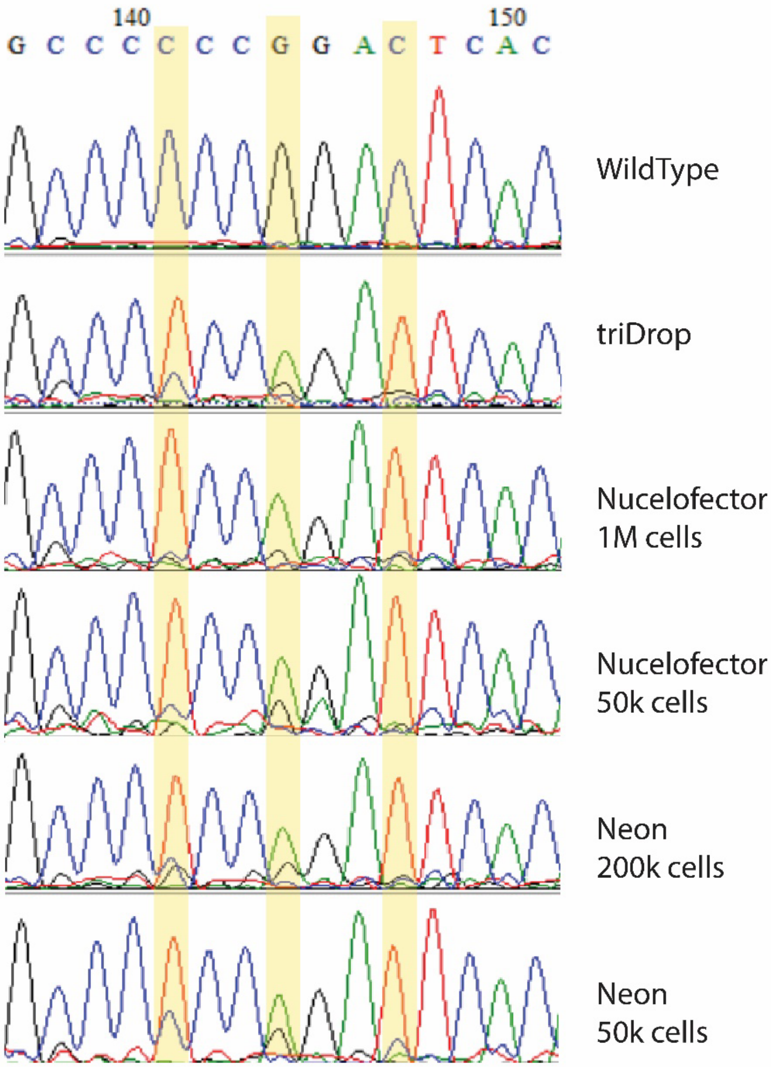


**Figure S5. Sanger Sequencing.** Wild type sequence and ssODN HDR template sequence with difference between the sequences highlighted. Chromatograms produced via Sanger for cells electroporated with all three EP systems using varying amounts of cells per reaction. Chromatograms are aligned using snap gene and = the yellow highlight is to show the difference between wild type and knock-in sequence.

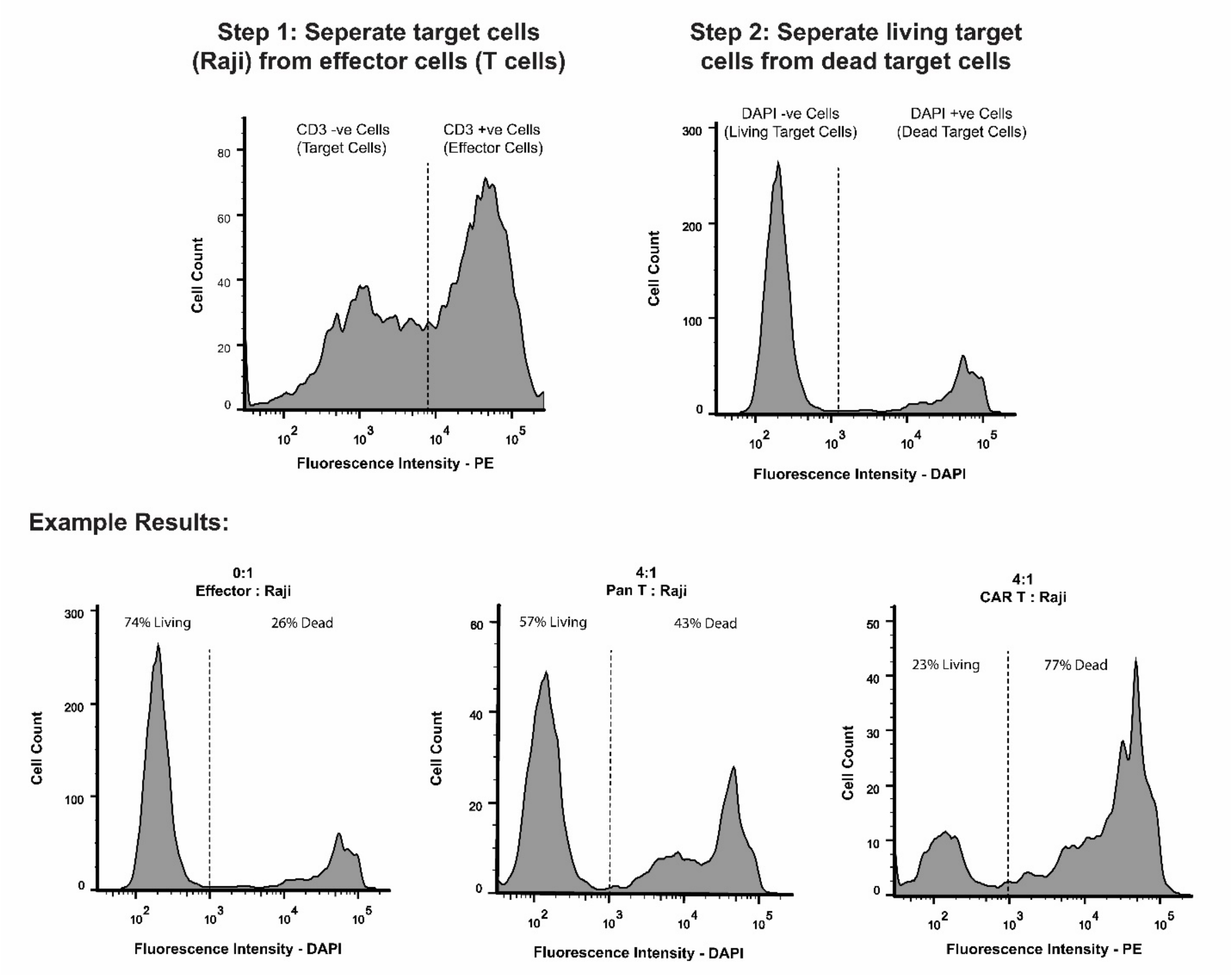


**Figure S6. CAR Killing-Assay Flow Cytometry Pipeline.** Flow cytometry gating pipeline for target cell killing analysis. Cells are stained with CD3-PE (Phycoerythrin) antibody and DAPI prior to flow cytometry. Histogram showing PE fluorescence intensity is generated for all lymphocytes showing two peaks allowing for effector and target cells to be differentiated. CD3 negative target cells (i.e. PE negative) are isolated and a new histogram is generated showing DAPI fluorescence intensity allowing for the differentiation of living and dead cells. Sample live-dead plots are shown for control Raji cells, and Raji cells cultured for 24 hours at a 4:1 ratio with activated pan T cells and triDrop engineered CAR T cells.
